# High-throughput discovery of peptide activators of a bacterial sensor kinase

**DOI:** 10.1101/2021.06.01.446581

**Authors:** Kathryn R. Brink, Andrew M. Mu, Ky V. Hoang, Ken Groszman, John S. Gunn, Jeffrey J. Tabor

## Abstract

Bacteria use two-component system (TCS) signaling pathways to sense and respond to peptides involved in host defense, quorum sensing, and inter-bacterial warfare. However, little is known about the peptide-sensing capabilities of these TCSs. Here, we develop a high-throughput *E. coli* display method to characterize the effects of human antimicrobial peptides (AMPs) on the pathogenesis-regulating TCS PhoPQ of *Salmonella* Typhimurium. We find that PhoPQ senses AMPs comprising diverse sequences, structures, and biological functions. Using thousands of AMP variants, we identify sub-domains and biophysical features responsible for PhoPQ activation. We show that most of the newfound activators induce PhoPQ in *S*. Typhimurium, suggesting a role in virulence regulation. Finally, we find that PhoPQ homologs from *Klebsiella pneumoniae* and extraintestinal pathogenic *E. coli*, which occupy different *in vivo* niches, exhibit distinct AMP response profiles. Our high-throughput method enables new insights into the specificities, mechanisms, and evolutionary dynamics of TCS-mediated peptide sensing in bacteria.

## Main Text

Peptide sensing plays an important role in bacterial quorum sensing, survival, and virulence. Bacteria use peptides to communicate with related strains^1,2^ and have evolved pathways to sense and respond to antimicrobial peptides (AMPs) involved in host defense^3,4^ and interbacterial warfare^5,6^. Peptide sensing is typically mediated by two-component system (TCS) signaling pathways. In a peptide-sensing TCS, peptides activate a membrane-bound sensor histidine kinase (SK), which autophosphorylates and transfers this phosphate group to its cognate cytoplasmic response regulator (RR). In pathogens, this RR often activates virulence pathways that enable these organisms to survive and replicate within the host environment^7^.

PhoPQ is a peptide-sensing TCS that regulates virulence in Gram-negative enterobacterial pathogens. This TCS is best studied in *Salmonella* Typhimurium. Typically *S*. Typhimurium bacteria enter the body through ingestion of contaminated food or water^8^. Once inside, they infect intestinal epithelial cells, cross the epithelial barrier, and are endocytosed by immune cells, including macrophages, where they survive and replicate intracellularly in a *Salmonella*-containing vacuole^8,9^. Here, the membrane-bound sensor histidine kinase PhoQ detects antimicrobial peptide (AMPs) produced as part of the human innate immune response, as well as acidic pH. PhoQ can also sense low divalent cation concentrations, though this stimulus is likely less important *in vivo*^10^. When activated, PhoQ phosphorylates the response regulator PhoP. Phosphorylated PhoP binds to genomic target promoters and directly or indirectly modulates transcription of more than 100 genes, including those involved in AMP resistance and virulence pathways that are essential for infection^11^.

Despite decades of investigation, little is known about the AMP-sensing capabilities of PhoQ. Previous studies of peptide-TCS interactions have largely relied on chemical peptide synthesis, which is costly and limited to short, linear peptides in high-throughput applications^12–15^. By contrast, host AMPs often contain disulfide bonds^16^ and can extend to tens of amino acids in length^17^. Due to these limitations, studies of AMP-PhoQ interactions have examined fewer than nine peptides each, most of which shared similar biochemical and biophysical properties^10,18–22^. These studies suggest that cationic AMPs bind to an acidic patch within the PhoQ periplasmic domain^10^, and that the extent of positive net charge and hydrophobicity are positively correlated with PhoQ activation^20^. Most of the 16 known PhoQ-activating AMPs, including the only known human activator cathelicidin LL-37 and its mouse ortholog cathelicidin-related antimicrobial peptide (CRAMP), are cationic and amphipathic^10,19–22^. Previously-described PhoQ activators are also predominantly α-helical^10,20,21^, with the notable exception of porcine protegrin-1, which adopts a β-sheet conformation^22^. However, positive charge and hydrophobicity alone are insufficient for PhoQ activation: β-sheet-structured α-defensins, such as the human AMP HNP-1, do not activate the sensor, despite being cationic and amphipathic^21,23^. These conflicting results and the small sample size of AMPs studied to date make it difficult to determine whether a given human AMP is likely to activate PhoQ. As a result, knowledge of human AMP-PhoQ interactions is limited to LL-37 and defensin HNP-1, the two AMPs that have been tested empirically.

In an effort to develop novel antimicrobial peptides, Davies and colleagues recently developed <u>s</u>urface-<u>l</u>ocalized <u>a</u>ntimicrobial displa<u>y</u> (SLAY), a multiplexed screening approach for determining the impact of peptides on bacterial viability^24^. In SLAY, *E. coli* are engineered to express a fusion protein that tethers peptides to the outer membrane (OM). The tether is of sufficient length that displayed peptides can penetrate the OM and interact with the periplasm and inner membrane (IM), yet short enough that they can only interact with the cell from which they are expressed. This *cis* activity enables peptide sequence to be directly linked to phenotype (i.e. growth) via <u>n</u>ext <u>g</u>eneration DNA <u>s</u>equencing (NGS). SLAY also permits the formation of disulfide bonds to enable surface display of AMPs with more complex structures, such as defensin HNP-1^24^. The Davies group used NGS to quantify differences in growth rate for 800,000 peptide-displaying strains in a mixed culture, identifying nearly 8,000 potential new AMPs^24^. Despite its demonstrated utility for AMP development, SLAY has not been utilized to characterize peptide-TCS interactions.

Here, we combine SLAY with heterologous TCS expression and sort-seq, a high-throughput fluorescent reporter gene expression assay based on fluorescence-activated cell sorting (FACS) with NGS, to measure the response of *S*. Typhimurium PhoPQ to 117 human AMPs and thousands of variants of those AMPs in laboratory *E. coli*. We name this method SLAY-TCS. Using SLAY-TCS, we identify 13 human AMP activators of PhoQ, evaluate the effects of subdomains of human AMP activators with combined α+β structures on PhoQ, and identify peptide biochemical and biophysical properties associated with PhoQ activation. We go on to validate these newfound activators using chemical and recombinant peptide synthesis and PhoPQ assays in *S*. Typhimurium. Finally, we use a simplified SLAY-TCS protocol wherein sort-seq is replaced with flow cytometry to discover that PhoPQ orthologs from extraintestinal pathogenic *E. coli* (ExPEC) and *Klebsiella pneumoniae* have evolved altered AMP sensing specificities relative to *S*. Typhimurium PhoPQ. In contrast to the previous literature, which has considered cationic AMPs to be interchangeable in their ability to activate this sensor^25,26^, our work demonstrates that PhoQ senses select human AMPs in a specific manner. Our results suggest a role for evolutionary adaptation in AMP sensing specificity and provide the first method for high-throughput characterization of peptide-TCS interactions.

## Results

### Porting *S*. Typhimurium PhoPQ into *E. coli*

To enable high-throughput screening of AMP-PhoQ interactions, we engineered an *E. coli* reporter strain of *S*. Typhimurium PhoPQ activity (**Figure 1a-b**). First, we constructed a genomic integration cassette wherein *S*. Typhimurium *phoP* and *phoQ* are expressed from constitutive promoters and other well-characterized gene regulatory elements (**Figure 1b, Supplementary Figure 1d**). We integrated the resulting engineered PhoPQ expression cassette into the genome of *E. coli* BW30007, which lacks the *E. coli phoPQ* ortholog^27^, resulting in strain KB1. Next, we developed a plasmid-based transcriptional reporter of PhoPQ activity. When overexpressed, PhoP can activate target promoters independently of PhoQ^28^. Thus, we constructed five plasmids, each containing an anhydrotetracycline (aTc)-inducible *phoP* and a different *S*. Typhimurium PhoP-activated promoter driving expression of a superfolder GFP (*sfgfp*)^29^ fluorescent reporter gene (**Supplementary Figure 1a**). We transformed these plasmids into BW30007 and measured sfGFP fluorescence in response to varying levels of aTc (**Supplementary Figure 2**). We found that the *virK* promoter (P_*virK*_) yields the largest fold activation in response to aTc exposure (**Supplementary Figure 2**). Thus, we selected P_*virK*_ as a reporter of PhoPQ activity for our subsequent experiments.

**Figure 1.**
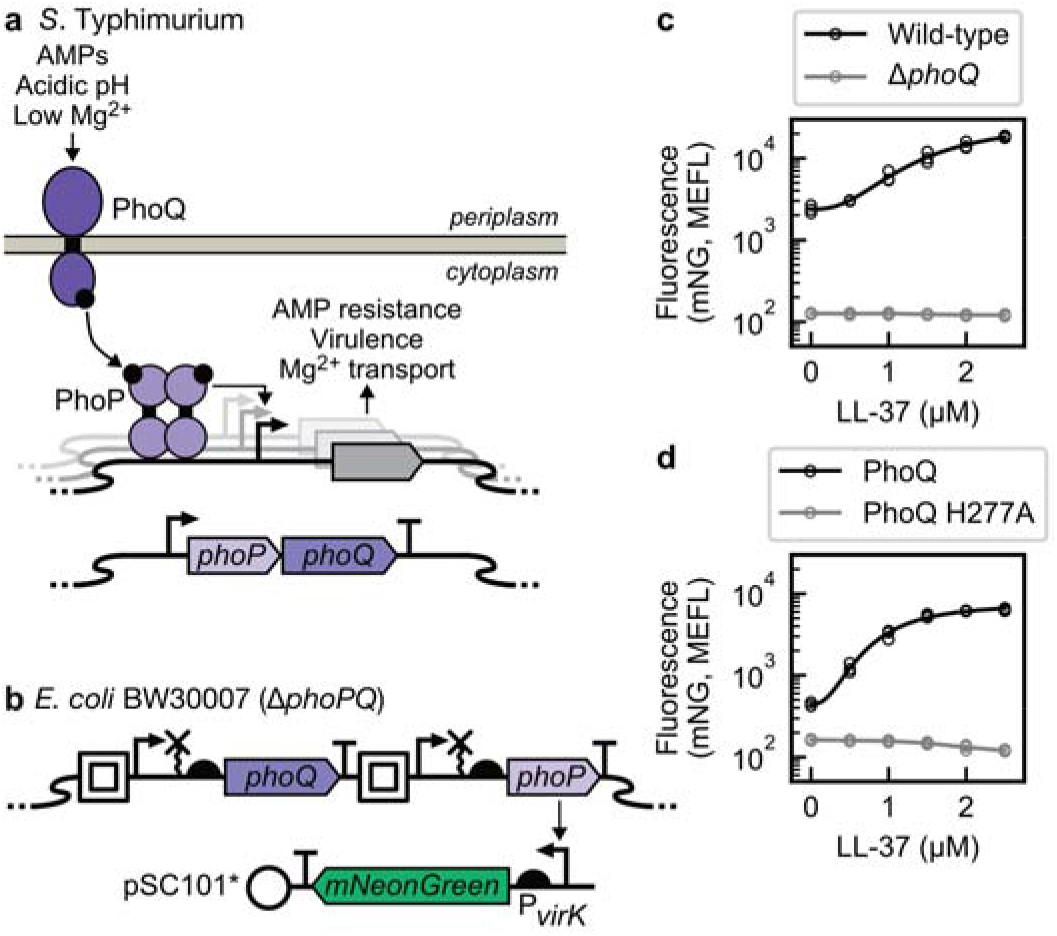
Porting *S*. Typhimurium PhoPQ into *E. coli*. (**a**) *S*. Typhimurium PhoPQ natively controls many physiological functions. (**b**) Genetic diagram of KB1, in which *S*. Typhimurium PhoPQ has been ported to laboratory *E. coli*, and of pKB233, the P*_virK_* fluorescent reporter plasmid (full genetic schematics in **Supplementary Figure 1**). (**c, d**) PhoPQ induction with LL-37 in *S*. Typhimurium (**c**) and engineered *E. coli* KB1 (**d**) strains containing pKB233, as measured by flow cytometry. Data was collected over *n* = 3 separate days, with results from each day shown as a separate marker.

Next, we constructed a plasmid wherein P*_virK_* controls expression of an *mNeonGreen* (*mNG*) fluorescent reporter gene, which produces a fluorescent protein that is brighter than sfGFP. We transformed this plasmid into both *S*. Typhimurium and KB1 to measure PhoPQ activation by the positive control AMP LL-37. We observed that this P*_virK_-mNG* reporter system is activated in a similar manner in both organisms (**Figure 1c-d**), with half-maximal activation occurring at 2.01 ± 0.43 and 1.16 ± 0.10 μM LL-37, respectively. We also measured LL-37 response in two negative control strains: an *S*. Typhimurium strain lacking *phoQ* and a KB1 derivative in which the catalytic histidine of PhoQ was mutated to an alanine (KB1 PhoQ H277A). Neither negative control strain responds to LL-37, demonstrating that *S*. Typhimurium PhoPQ specifically senses this AMP in both its native context and in *E. coli* KB1. These results demonstrate that we have recapitulated the function of *S*. Typhimurium PhoPQ in *E. coli* strain KB1.

### PhoPQ senses surface-displayed peptides in *E. coli*

Next, we validated that PhoPQ can detect AMPs displayed on the surface of KB1. In SLAY, peptides of interest are expressed at the C-terminus of a fusion protein consisting of the Lpp signal peptide, an OmpA transmembrane domain, and a flexible tether^24^. Here, we constructed a series of 13 displayed peptide expression plasmids adapted from our P*_virK_-mNG* reporter plasmid. In addition to P*_virK_-mNG*, each of these plasmids encodes a SLAY fusion protein under the control of an isopropyl-β-D-thiogalactoside (IPTG)-inducible promoter (P_tac_) (**Figure 2a**, **Supplementary Figure 1c**). P_tac_ is tightly repressed, enabling us to avoid potential toxicity arising from leaky AMP expression in the absence of inducer. Using this platform, we measured the PhoPQ response to nine positive control peptides that had been shown to activate PhoPQ in previous studies (**Figure 2c-d**). We included cecropin P1, which had previously been displayed with SLAY^24^, as a tenth putative activator, as a *phoQ* mutant strain had previously exhibited increased susceptibility to this peptide^30^ (**Figure 2c-d**). We also measured two negative control peptides: human defensin HNP-1^23^ (**Figure 2b, d**), which had previously been shown not to activate PhoPQ, and the scrambled, inert peptide NC^19^, which had previously been shown to be non-toxic to *S*. Typhimurium *phoP* and *phoQ* mutant strains. Finally, we constructed a no-peptide control plasmid encoding a stop codon instead of a peptide following the OmpA transmembrane domain. This plasmid enables us to quantify the baseline level of fluorescence from our P*_virK_-mNG* reporter in the absence of any displayed peptide in order to calculate a fold change of PhoPQ activation for each AMP.

**Figure 2.**
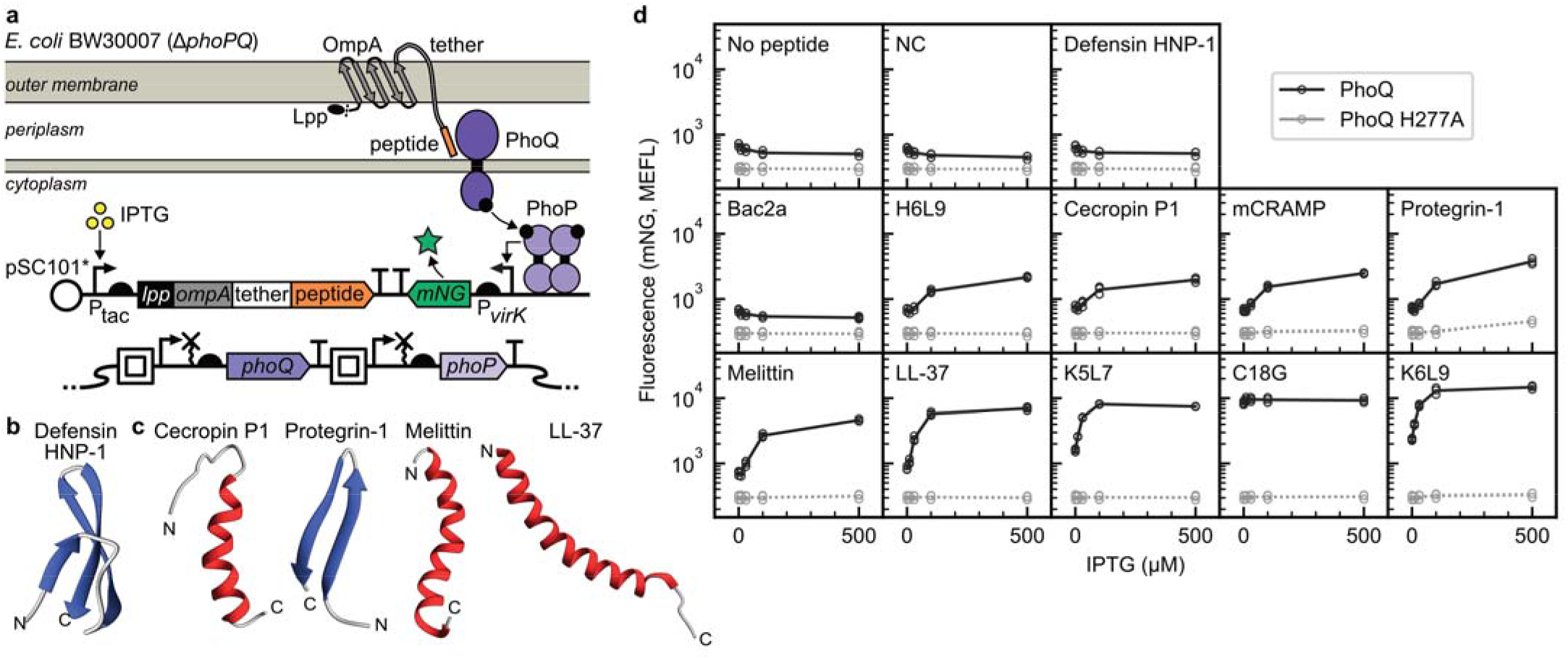
Surface-displayed AMPs activate PhoPQ in *E. coli*. (**a**) Schematic of peptide surface display and genetic diagram of peptide surface display and mNeonGreen fluorescent reporter plasmids (full genetic schematics in **Supplementary Figure 1**). (**b**) Structure of the only known non-activating human AMP (PDB ID – HNP-1: 3GNY). (**c**) All available structures of known PhoQ-activating AMPs (PDB ID – cecropin P1: 2N92, protegrin-1: 1PG1, melittin: 6DST, LL-37: 2K6O). (**d**) PhoPQ activation in KB1 in response to IPTG induction of surface-displayed AMPs, as measured by flow cytometry. Data was collected over *n* = 3 separate days, with results from each day shown as a separate marker.

As expected, the no-peptide control, human defensin HNP-1, and NC do not activate PhoPQ when displayed in KB1 (**Figure 2d**). On the other hand, nine out of the 10 positive control peptides induce mNG expression between 8.3- and 71-fold at maximum IPTG induction (500 μM) (**Figure 2d**). Notably, protegrin-1 activates PhoPQ when surface-displayed, indicating that SLAY-TCS is compatible with AMPs containing disulfide bonds. Of the positive control peptides, only Bac2a does not activate PhoPQ. In the previous study where Bac2a was reported to activate PhoPQ in *S*. Typhimurium, it was amidated on its C terminus^19^, a modification that is not possible using SLAY. Our data suggest that this C-terminal amidation, which also increases the net charge of the peptide by +1, may be important for Bac2a sensing by PhoPQ. To validate that activation of PhoPQ by surface-displayed AMPs is dependent on PhoQ kinase activity, we repeated this assay in KB1 PhoQ H277A. As expected, we observed no PhoPQ activation for any peptide in the PhoQ H277A strain (**Figure 2d**). We did observe a slight increase in mNG fluorescence at very high levels of protegrin-1 expression (**Figure 2d**). However, this effect is likely due to an AMP-mediated growth slowdown resulting in mNG accumulation (**Supplementary Figure 3**). We conclude that SLAY-TCS enables faithful characterization of the response of PhoPQ to surface-displayed peptides, including those containing disulfide bonds.

### PhoQ senses diverse human AMPs

To characterize the response of PhoQ to most human AMPs, we first identified 133 human AMPs in the Antimicrobial Peptide Database (APD3)^17^. From this set, we excluded 16 (12%) exceeding 104 amino acids in length due to size restrictions of commercial oligonucleotide synthesis (**Supplementary Figure 4a**, **Methods**). The remaining set of 117 AMPs vary in charge, length, hydrophobicity, and secondary structure (**Supplementary Figure 4a-d**, **Supplementary Table 1**), and comprise 11 clusters of high sequence similarity (**Supplementary Figure 4e**, **Supplementary Table 1**). These clusters include families of mature AMPs processed from a single precursor peptide (e.g. products of cathelicidin, human defensin 5, and β-amyloid) and peptides with high sequence similarity that are produced from distinct genes (e.g. CXCL1, CXCL2, and CXCL3).

Next, we constructed a human AMP plasmid expression library wherein each plasmid encodes P*_virK_-mNG* and a surface-displayed human AMP under control of P_tac_, as in our control studies (**Methods**). We transformed this plasmid library into KB1. Using Illumina NGS, we validated that sequences encoding all 117 human AMPs, along with no-peptide controls, are present in the KB1 human AMP library. We also identified variants of human AMPs that likely resulted from errors in oligo synthesis or plasmid assembly. As expected, these variants generally had lower representation in the library than the human AMPs (**Supplementary Figure 5b**).

Next, we utilized sort-seq to characterize the effect of each peptide on PhoPQ activity. First, we grew the KB1 human AMP library in the presence of 500 μM IPTG, which induces sufficient expression of positive control AMPs to strongly activate PhoPQ (**Figure 2d**). Then, we used FACS to sort these bacteria into eight logarithmically-spaced mNG fluorescence bins (**Figure 3b**, **Supplementary Table 2**). The bins were designed to span the fluorescence distributions of negative (no peptide) and positive (C18G) PhoQ activity control strains measured on the same day (**Supplementary Figure 5a**). Induction of the peptide library resulted in two distinct mNG fluorescence peaks corresponding to PhoPQ activity levels of our positive and negative controls (**Supplementary Figure 5a**). This result suggested that some peptides in the library strongly activate PhoQ. IPTG treatment had little effect on bacterial growth, suggesting that peptide toxicity did not substantially influence our measured fluorescence values (**Supplementary Figure 5b**). Next, we used Illumina NGS to sequence the peptide-encoding DNA regions from bacteria in each bin and estimated the mean mNG fluorescence level of each strain using the relative abundances of peptide-encoding sequences across fluorescence bins. We divided these estimated fluorescence values by the estimated mean fluorescence of our no-peptide control to calculate the fold activation of PhoPQ in response to each peptide in the library. These analyses revealed that LL-37 activates mNG expression 5.2-fold, whereas HNP-1 does not activate mNG expression (**Figure 3c**), consistent with previous work using purified peptides.

**Figure 3.**
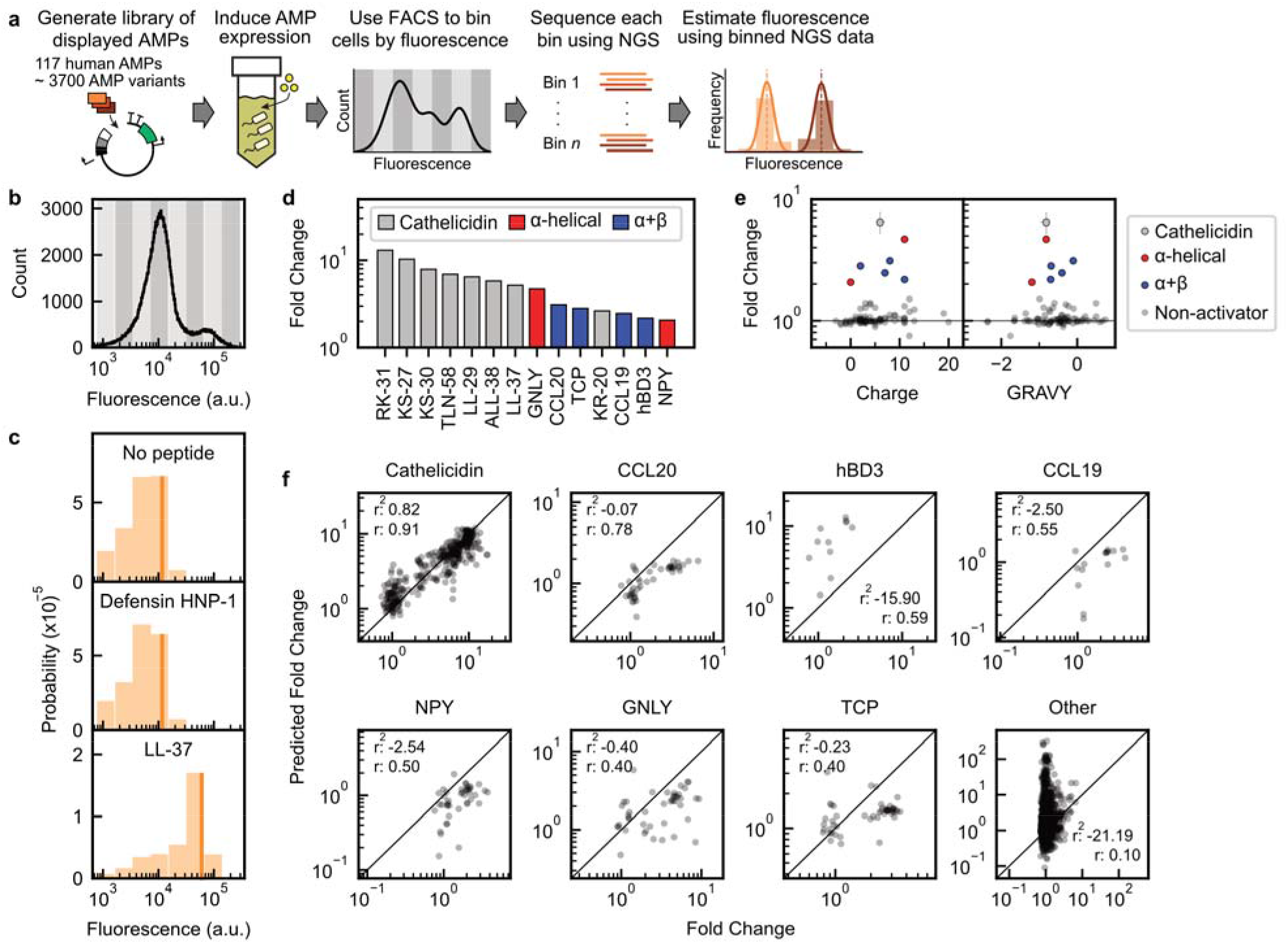
Discovery and characterization of PhoPQ-activating human AMPs. (**a**) SLAY-TCS workflow. (**b**) Sort gate strategy for sort-seq, with sorting bins represented by alternating light and dark grey bars (see **Supplementary Table 2** for bin edge fluorescence values). Histogram represents the results from a single biological replicate (*n* = 1). (**c**) Calculated fluorescence distributions of no peptide, negative (defensin HNP-1), and positive (LL-37) control strains included in the library. Vertical lines indicate estimated mean fluorescence values for each control. (**d**) Sort-seq fold activation relative to no peptide control strains for all human AMP activators with > 2-fold activation. (**e**) Net charge and hydrophobicity, as measured by grand average of hydropathicity (GRAVY), for all human AMPs included in the screen. Human AMPs belonging to the same cluster (**Supplementary Table 1**) are represented by a single data point with error bars indicating the median and 25^th^ to 75^th^ percentile ranges for that cluster. (**f**) Performance of the cathelicidin sparse robust linear model on peptide variants of human AMP activators of PhoPQ (cathelicidin, CCL20, hBD3, CCL19, NPY, GNLY, TCP) and on peptides variants of non-activating human AMPs (Other). Coefficient of determination (*r*^2^) and Pearson correlation coefficient (*r*) values were calculated for log_10_-transformed fold change and predicted fold change values.

In addition to LL-37, we identified 13 AMPs that activate PhoPQ activity at least 2-fold in our sort-seq screen (**Figure 3d**). Seven of these AMPs are derived from cathelicidin, the same parent protein as LL-37, and exhibit substantial sequence identity to LL-37 (**Supplementary Figure 5c**, **Supplementary Table 1**). Only one cathelicidin-derived AMP, LL-23, failed to activate PhoPQ by 2-fold in our assay (**Supplementary Figure 5c**, **Supplementary Table 1**). The remaining six activators are the chemokines CCL19 and CCL20, granulysin (GNLY), human β-defensin 3 (hBD3), thrombin-derived C-terminal peptide (TCP), and neuropeptide Y (NPY), which are unrelated to cathelicidin. We validated that each of these non-cathelicidin peptides specifically activates PhoQ when surface-displayed in KB1 in single-strain cultures (**Supplementary Figure 5d**).

The 6 non-cathelicidin PhoQ activators exhibit diverse sequences, structures, biochemical properties, and physiological functions. GNLY and NPY exhibit predominantly α-helical folds (**Supplementary Figure 5e**). The remaining four (hBD3, CCL19, CCL20, TCP) adopt combined α+β structures (**Figure 4a**) and are the first reported PhoQ activators to exhibit this structure. These AMPs also vary in their net charge and hydrophobicity (**Figure 3e**, **Supplementary Table 1**). In fact, NPY has a net neutral charge in our assay and is the first reported non-cationic PhoPQ activator. In addition to their antimicrobial activity, these AMPs play important roles in immunomodulation and other biological process across diverse sets of human tissues (**Supplementary Figure 6a**)^31–41^. GNLY and CCL20, are expressed in *S*. Typhimurium-exposed human peripheral blood mononuclear cells (PBMCs) (**Supplementary Figure 6b**), which include macrophages, an important infection niche for *S*. Typhimurium^42^. To determine whether macrophages are responsible for the GNLY and CCL20 expression observed in PBMCs, we exposed macrophages derived from a human monocytic cell line (THP-1) and primary human monocytes to *S*. Typhimurium. We did not detect GNLY or CCL20 in these cells (**Supplementary Figure 6c-d**), suggesting that PhoQ activation by these AMPs may occur via other cell types. This finding is consistent with previous work that did not detect AMP-mediated PhoQ activation in macrophages *in vitro*, but did observe AMP-mediated PhoQ activation *in vivo* in the mouse gut lumen^21^. Together, these results reveal that diverse human AMPs activate PhoPQ and support previous work suggesting that PhoPQ activation may occur prior to *S*. Typhimurium entry into macrophages.

**Figure 4.**
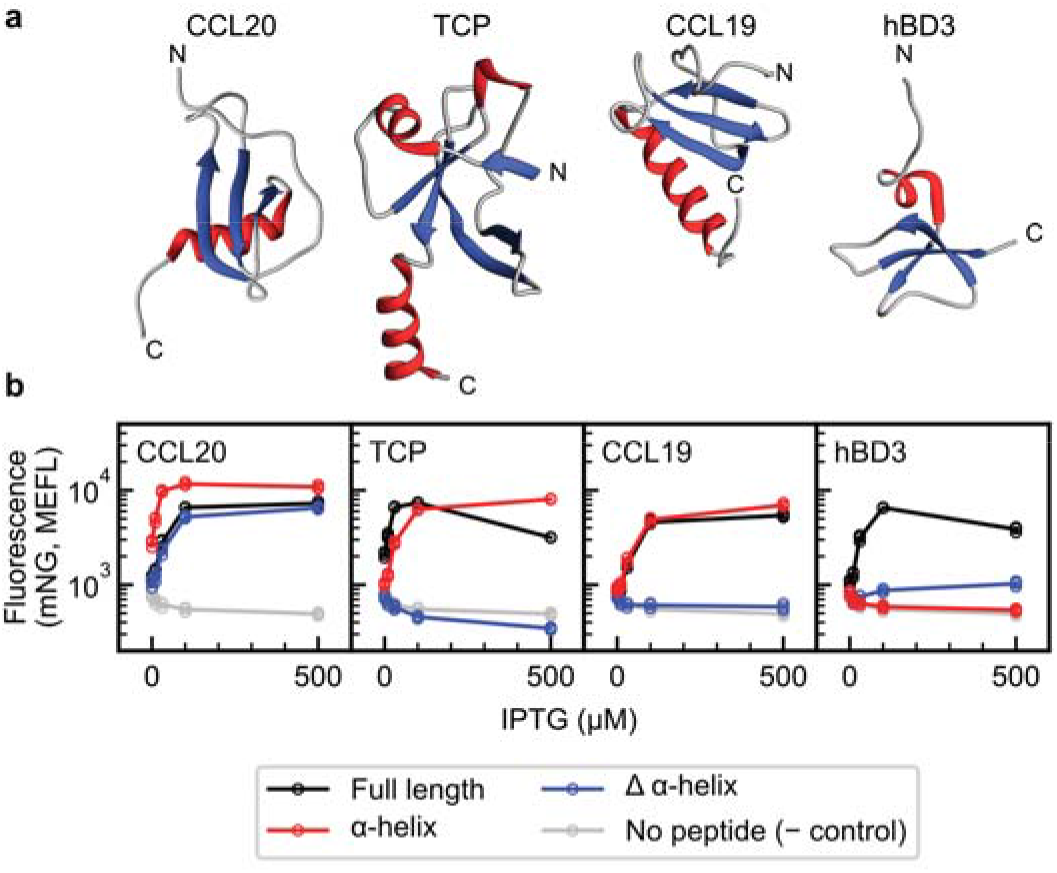
Validation of SLAY-TCS hits and follow-up truncation studies. (**a**) Structures of α+β human AMP activators. (PDB ID – CCL20: 2JYO, TCP: adapted from 3U69, CCL19: 2MP1, hBD3: 1KJ6). (**b**) PhoPQ activation in KB1 in response to IPTG induction of surface-displayed truncation mutants (sequences in **Supplementary Figure 7**) as measured by flow cytometry. Data was collected over *n* = 3 separate days, with results from each day shown as a separate marker.

### PhoQ activators have diverse structure, charge, and hydrophobicity

Due to errors inherent in the oligonucleotide synthesis process, our surface-displayed AMP library contains at least 3,680 AMP variants (**Data availability**). Among these are at least 410 truncation and mutation variants of cathelicidin. Cathelicidin variants constitute the largest group of related peptides in our dataset and vary substantially in their ability to activate PhoPQ. Thus, we utilized these variants to identify cathelicidin regions and peptide biochemical and biophysical properties most important for PhoPQ activation.

To determine which cathelicidin regions are involved in PhoPQ activation, we compared our sort-seq data for 34 peptides: the 9 cathelicidin-derived AMPs that we designed into our library and the 25 cathelicidin truncation mutants ≥ 3 amino acids in length present due to errors (**Supplementary Figure 5c**). We found that the N-terminal region of TLN-58, the longest cathelicidin derivative in our library, does not independently activate PhoPQ (**Supplementary Figure 5c**). Instead, the strongest activators of PhoPQ are peptides in which the N-terminal residue corresponds to amino acid 140 or 141 of the CAMP gene product (**Supplementary Figure 5c**). This group includes the strongest human AMP activator in our library, RK-31 (**Figure 3d**), as well as a minimal PhoPQ-activating fragment, KSK (**Supplementary Figure 5c**). Thus, PhoPQ responds differently to different cathelicidin-derived peptides, despite their high sequence similarity.

To identify peptide features involved in PhoQ activation, we calculated 1,284 biochemical and biophysical properties from the primary amino acid sequences of each of the 410 cathelicidin variants (**Supplementary Note 2**). These include bulk features such as net charge and hydrophobicity, and detailed features arising from the order of amino acid residues along the peptide backbone (**Supplementary Table 3**). We then fit the cathelicidin variant sort-seq data to a sparse robust linear model using a training set of 308 peptides and validated model performance using a test set of 102 peptides (**Supplementary Note 3**). This machine learning approach narrows large feature sets into smaller feature sets that are highly predictive of the model output, in this case fold change of activation of PhoPQ^43^. Our model identified 14 properties that, when linearly combined, are highly predictive of PhoPQ activation for both the training and test sets of cathelicidin variants (**Supplementary Table 4**, **Supplementary Figure 5f**). Consistent with previous work, we find that positive net charge is correlated with PhoPQ activation (**Supplementary Table 4**). Surprisingly, GRAVY, a measure of bulk hydrophobicity, is not among the properties selected by our model (**Supplementary Table 4**). Instead, we identified several properties that involve correlations in hydrophobicity between residues separated by specified distances along the peptide backbone. Specifically, in PhoPQ-activating peptides, residues separated by a distance of 7 amino acids exhibit similar hydrophobic character (**Supplementary Table 4**). Furthermore, residues separated by 8 or 9 amino acids tend to exhibit similarities in hydrophobicity, hydrophilicity, and side chain mass (**Supplementary Table 4**). Finally, residues separated by 10 amino acids exhibit opposite hydrophobic character in PhoPQ activators (**Supplementary Table 4**). These findings indicate that the order and distribution of hydrophobic residues along a peptide are more important for PhoPQ activation than bulk hydrophobicity alone.

To explore the generality of these rules, we next evaluated whether our cathelicidin-trained model can predict PhoPQ activation by other classes of AMPs. To this end, we first applied our model to variants of the six non-cathelicidin activators that we identified in our sort-seq screen. The model successfully predicts trends in activation for most classes of AMP activators, as reflected by their Pearson correlation coefficients (*r* = 0.40 to 0.78) (**Figure 3f**). The model performs more poorly for variants of non-activating human AMPs (*r* = 0.10, Other in **Figure 3f**). Thus, while our model is generally effective at predicting trends in PhoPQ activation based on peptide sequence, additional properties involved in PhoPQ activation likely remain to be discovered.

### Helical and □-sheet regions of human AMPs activate PhoPQ

CCL19, CCL20, hBD3, and GNLY exhibit a combined α+β structure (**Figure 4a**), which has not been previously reported for any PhoQ-activating AMP. Given that nearly all previously-known PhoQ-activating AMPs are α-helical, we hypothesized that the α-helical regions of these activators may be responsible for PhoQ activation. To examine this hypothesis, we constructed sets of two truncation mutants for each AMP, one containing only the α-helix fragment of the AMP and a second lacking this α-helical sequence (**Supplementary Figure 7**). TCP contains several helical regions (**Figure 4a**). Thus, we chose to examine the longest α-helical region of TCP in our analysis, its C-terminal α-helix.

As expected, we found that the α-helix regions of CCL19 and TCP are both necessary and sufficient for PhoPQ activation (**Figure 4b**). Surprisingly, both the C-terminal α-helix and N-terminal β-sheet regions of CCL20 independently activate PhoPQ. Finally, neither hBD3 mutant activates PhoPQ. Truncating hBD3 between its α-helix and β-sheet regions disrupted a disulfide bond. Disulfide bonds are required for chemoattraction by hBD3^31^ and may be required for hBD3 activation of PhoPQ. We conclude that while the α-helical regions of AMPs with α+β structures are generally sufficient for PhoPQ activation, β-sheet regions can also play an important role in activating this sensor.

### Non-cathelicidin AMPs activate PhoPQ in *S*. Typhimurium

Next, we examined whether the non-cathelicidin AMPs that activate PhoPQ when displayed on the surface of *E. coli* similarly activate PhoPQ when delivered exogenously to *S*. Typhimurium. To this end, we sourced full-length CCL19, GNLY, CCL20, NPY, and hBD3 from commercial suppliers. TCP was not available commercially. Instead, we had the active C-terminal α-helix fragment custom synthesized (**Figure 5c**). We used the *S*. Typhimurium P*_virK_-mNG* reporter strain that we developed previously (**Figure 1c, Supplementary Figure 1b**) to assess these AMPs for PhoQ activation in the pathogen. We found that all full-length AMPs except CCL19 activate PhoPQ by flow cytometry (**Figure 5a-b**). Because the C-terminal α-helix fragment of TCP was custom synthesized, we were able to obtain sufficient peptide to measure concentration-dependent activation of PhoPQ. We observed half-maximal activation of PhoPQ by TCP C-terminal α-helix at 1.24 ± 0.10 μM (**Figure 5d**), which is comparable to the half-maximal activation for LL-37 (**Figure 1d**). Finally, all of these responses are abolished in an *S*. Typhimurium *phoQ* knockout strain, demonstrating that they arise due to PhoQ-mediated sensing. These experiments reveal that GNLY, CCL20, NPY, hBD3, and the TCP C-terminal α-helix fragment activate PhoPQ in *S*. Typhimurium.

**Figure 5.**
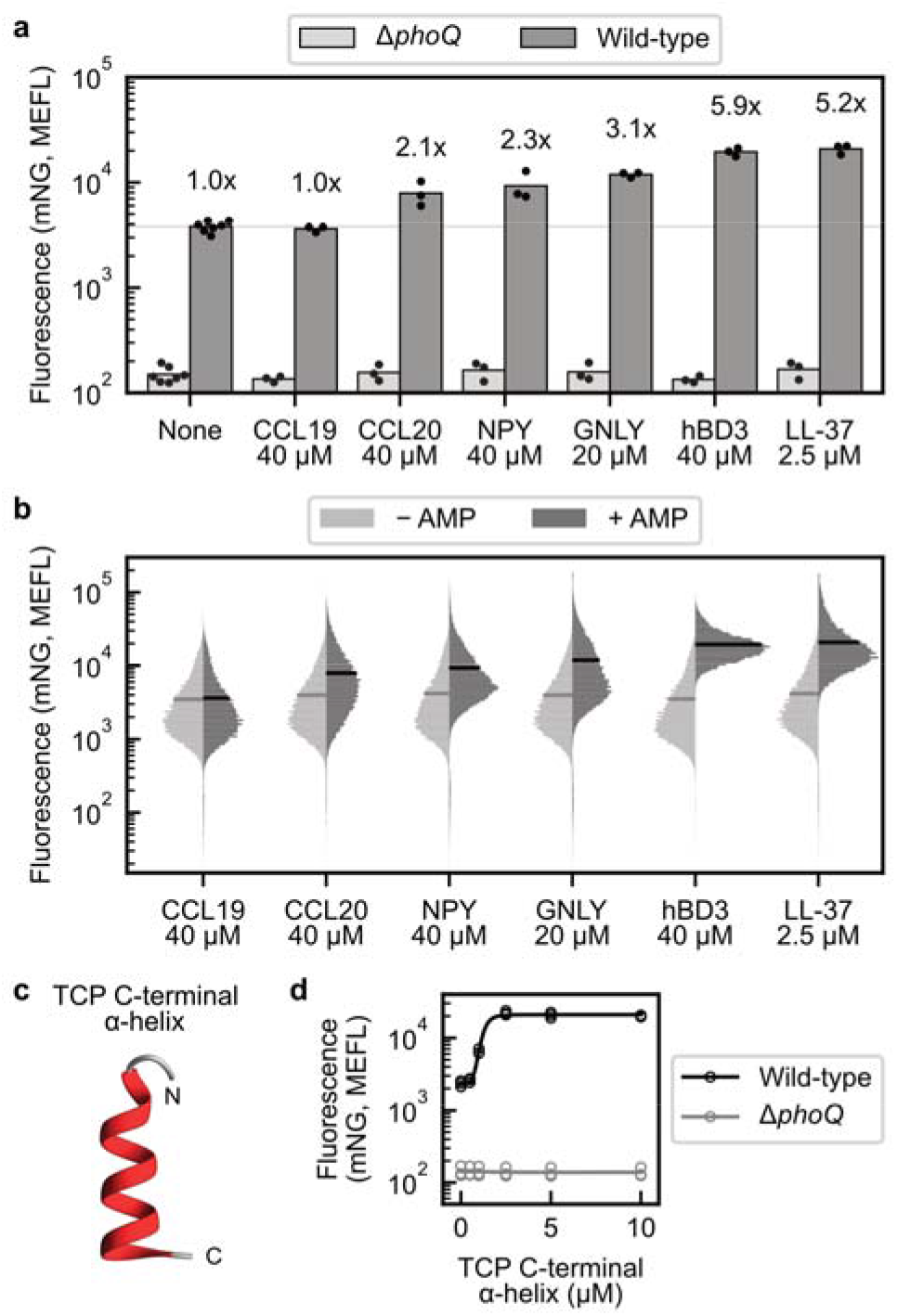
Non-cathelicidin AMPs activate PhoPQ in *S*. Typhimurium. (**a**) PhoPQ activation by full-length AMPs in pKB233-containing wild-type and *ΔphoQ S*. Typhimurium strains, as measured by flow cytometry. Data for each AMP was collected in *n* = 3 separate day replicates; controls where no AMP was added were collected alongside each AMP replicate, for a total of *n* = 7 separate day replicates. Each marker represents one biological replicate; bar heights correspond to the average of these replicates. Fold changes (above the bars) are the mean autofluorescence-subtracted fold changes between AMP-containing and control samples collected on the same day. (**b**) Flow cytometry histograms of the pKB233-containing wild-type *S*. Typhimurium strain in response to full-length AMPs. Each histogram is the combined fluorescence distribution for *n*□=□3 separate day experiments. Side-by-side histograms for control (– AMP) and AMP-containing (+AMP) samples were collected on the same day. (**c**) Structure of C-terminal α-helix fragment of TCP (adapted from PDB 3U69). (**d**) PhoPQ activation by TCP C-terminal α-helix fragment in pKB233-containing wild-type and Δ*phoQ S*. Typhimurium strains, as measured by flow cytometry. Data was collected over *n* = 3 separate days, with results from each day shown as a separate marker.

### PhoPQ orthologs exhibit altered AMP-sensing specificity

Although PhoPQ is highly conserved among Enterobacteriaceae, PhoQ orthologs exhibit substantial sequence variability, particularly in the periplasmic domain where AMP sensing is believed to occur. For example, PhoQ orthologs from ExPEC and *K. pneumoniae*, two enterobacterial species that commonly exhibit high antibiotic resistance, are 85% and 81% identical to *S*. Typhimurium PhoQ overall, yet their periplasmic domains are only 81% and 74% identical to that of *S*. Typhimurium PhoQ (**Supplementary Figure 8a**). Several non-conserved residues are located in the PhoQ acidic patch, an important region for AMP sensing^10^ (**Figure 6a**). We hypothesized that sequence variability in the acidic patch region could alter AMP sensing specificity, which might enable pathogens to sense AMPs that could confer them with an evolutionary advantage during infection.

**Figure 6.**
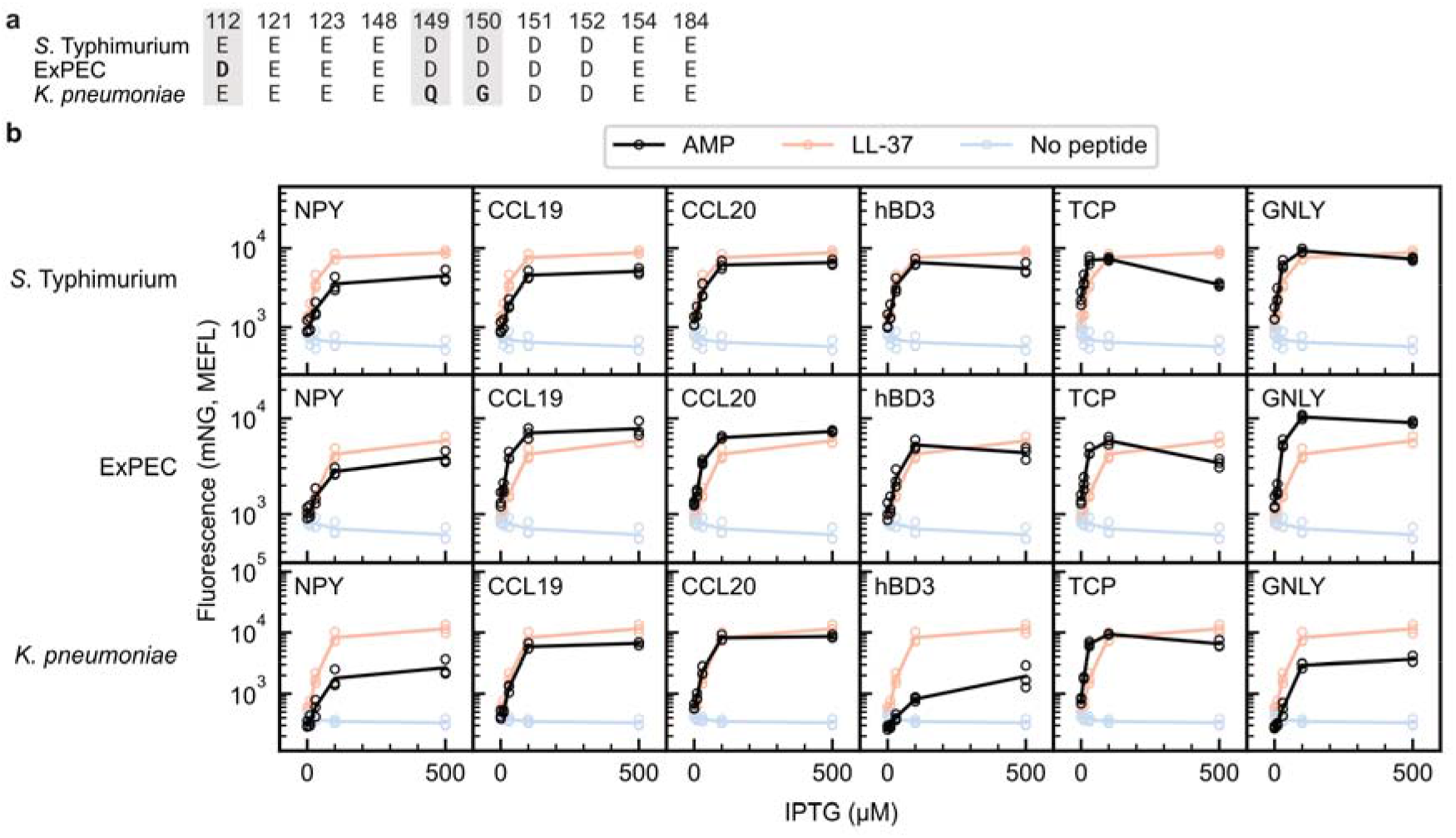
*S*. Typhimurium, ExPEC, and *K. pneumoniae* PhoPQ exhibit different sensitivities to surface-displayed AMPs. (**a**) *S*. Typhimurium, ExPEC, and *K. pneumoniae* PhoQ acidic patch residues. Residues are numbered relative to *S*. Typhimurium; variable residues across these orthologs are shaded and mutations in the acidic patch relative to *S*. Typhimurium are bolded. (b) PhoPQ activation in KB1, AM1, and AM2 in response to IPTG induction of surface-displayed AMPs as measured by flow cytometry. Data was collected over *n* = 3 separate days, with results from each day shown as a separate marker.

To test this hypothesis, we ported ExPEC and *K. pneumoniae* PhoPQ into *E. coli* BW30007 to generate strains AM1 and AM2, respectively. These orthologs are expressed from a similar engineered genetic context to *S*. Typhimurium PhoPQ in KB1 (**Supplementary Figure 1d**). Given that *S*. Typhimurium, ExPEC, and *K. pneumoniae* PhoP are closely related, we hypothesized that all orthologs would activate our *S*. Typhimiurium-derived P_*virK*_ reporter. To validate P_*viřK*_ activation, we transformed AM1 and AM2 with our P_*virK*_-*mNG* reporter plasmid and assessed their response to Mg^2+^ limitation (**Supplementary Figure 8b**). All three orthologs are activated in limiting Mg^2+^ conditions, with *K. pneumoniae* PhoPQ exhibiting a more modest response (3.8-fold between 10 μM and 10 mM Mg^2+^) than *S*. Typhimurium and ExPEC PhoPQ (12.2- and 37.9-fold, respectively) in our engineered *E. coli* context. These data confirm that we have functionally ported ExPEC and *K. pneumoniae* PhoPQ into *E. coli*.

To determine whether PhoPQ orthologs differ in their AMP-sensing specificities, we displayed LL-37 and the six non-cathelicidin human AMP activators along with a no peptide control in KB1, AM1, and AM2 (**Figure 6b**). We induced AMP expression with IPTG and measured PhoPQ activation in each strain using our P*_virK_-mNG* reporter. Despite its weaker Mg^2+^ response, *K. pneumoniae* PhoPQ exhibits a similar maximum response to displayed peptides as ExPEC and *S*. Typhimurium PhoPQ (**Supplementary Figure 8b**). This finding confirms that displayed peptides activate *K. pneumoniae* and ExPEC PhoPQ in *E. coli* and suggests that *K. pneumoniae* PhoPQ may be less sensitive to Mg^2+^ limitation than ExPEC and *S*. Typhimurium PhoPQ.

Finally, we found that while all three PhoPQ orthologs respond to all of the AMP activators, their response profiles differ. For example, while each ortholog responds strongly to CCL20, their responses to GNLY vary greatly (**Figure 6b**). In particular, ExPEC PhoPQ responds more strongly to GNLY than to LL-37, while *K. pneumoniae* PhoPQ exhibits a weaker response to this AMP. Similarly, *K. pneumoniae* PhoPQ responds more weakly to hBD3 than *S*. Typhimurium or ExPEC PhoPQ. Interestingly, these AMPs have varying tissue expression patterns within the human body. CCL20 is expressed across many tissues, whereas GNLY is primarily expressed in immune cells, such as T cells and B cells (**Supplementary Figure 6a**). hBD3 is expressed predominantly in oral cavity and esophageal tissues (**Supplementary Figure 6a**). These results show that PhoPQ-mediated AMP sensing differs between orthologs and present the possibility that AMP sensing specificity can evolve to increase pathogen fitness in different infection sites.

## Discussion

Peptides are an important class of TCS activators. However, the extent and specificity of peptide-SK interactions remains largely unknown. The major challenge has been a reliance on low-throughput peptide synthesis and purification techniques. By combining peptide surface display with TCS engineering, SLAY-TCS enabled us to perform the largest peptide-TCS interaction study to date by two orders of magnitude. Through this work, we discovered novel interactions between human AMPs and a key virulence-regulating TCS, used data from hundreds of peptides to identify peptide properties involved in sensor activation, and uncovered differences in specificity between sensor orthologs that suggest a role for evolutionary adaptation in AMP sensing. These findings not only deepen our understanding of the role of PhoPQ in host-pathogen interactions, they provide a proof-of-principle for an approach that can be extended to characterize a wide range of bacterial sensors.

Prior to this study, cathelicidin LL-37 was the only known human AMP activator of PhoPQ. Using SLAY-TCS, we discovered 13 additional human AMP activators of this TCS. These AMPs include the first PhoPQ activators with combined α+βstructure (CCL19, CCL20, GNLY, TCP), the first activator derived from the β-sheet region of a human AMP (CCL20 C-terminal fragment), and the first non-cationic activator (NPY). One previous study of peptide variants identified positive charge and hydrophobicity as features that positively correlate with PhoQ activation^20^. However, this study examined only eight peptides^20^. By applying machine learning to data collected on the PhoPQ response to hundreds of cathelicidin sequence variants, we built on the results of this previous study and validated that positive net charge is positively correlated with PhoQ activation^20^. In addition, our data suggests a new role for long-range correlations in hydrophobicity in peptides that activate PhoQ. Surprisingly, our machine learning model is also able to predict general trends in PhoQ activation for classes of AMPs with no homology to cathelicidin, suggesting that many of the properties that we identified could be generally important for peptide-mediated activation of this sensor. While other studies of ligand-receptor interactions have exhibited superior performance to our approach, these efforts have typically used binary classification schemes, employed more complex structural biophysical models, or relied on blackbox models with limited interpretability^44–46^. By contrast, our model relies only on peptide primary amino acid sequences and uses a highly interpretable algorithm to correlate peptide features with PhoPQ activation. Future studies with larger and more diverse peptide libraries could enable more powerful machine learning models that can identify complex, non-linear, and generalizable associations between peptide properties and PhoPQ activation. Specialized peptide libraries could also be designed to probe the importance of specific sequence, structural, biochemical, and biophysical characteristics.

The PhoPQ-activating AMPs that we discovered have diverse physiological functions. As chemokines, CCL19 and CCL20 regulate immune cell migration through binding to their cognate receptors CCR7 and CCR6, respectively^32,34,35^. CCL19 is produced constitutively and plays a role in adaptive immunity^32,34^, whereas CCL20 is upregulated in inflammation and is implicated in mucosal immunity in the gut^35^. GNLY and hBD3 are primarily known for their antimicrobial properties but have also been shown to exhibit immunomodulatory effects^36,37^. In fact, hBD3 may recruit some immune cells through interactions with CCR6^31^, the chemokine receptor for CCL20. Thrombin-derived C-terminal peptides play a role in wound healing and immunomodulation and can aggregate Gram-negative and Gram-positive bacteria, likely through interactions with lipopolysaccharide, lipid A, and lipoteichoic acid^38,39^. NPY is a widely expressed neurotransmitter, the most abundant neuropeptide in the brain, and an important immunomodulatory peptide in the gut^40^. Future studies will be required to validate PhoPQ activation by the AMPs that we identified in *in vitro* cell culture models and through *in vivo* studies of *S*. Typhimurium infection. Our findings regarding the expression of these AMPs in human tissues, PBMCs, and macrophages (**Supplementary Figure 6**) suggest that PhoQ activation may occur outside of the macrophage environment, in line with previous work demonstrating a role for AMP-mediated activation of PhoQ within the gut lumen^21^. Future *in vivo* studies could provide additional insights into PhoQ activation by AMPs prior to *S*. Typhimurium entry into macrophages.

SLAY-TCS results are extensible to TCSs in their native contexts. Five out of six of the non-cathelicidin AMPs that we identified activate PhoPQ when exogenously supplied to *S*. Typhimurium. One AMP, CCL19, did not activate PhoPQ in *S*. Typhimurium in our experiments. A possible explanation for this discrepancy is the presence of O-antigen, a decoration present on the outer membrane of *S*. Typhimurium but absent from our *E. coli* strain^47^. O-antigen may have inhibited CCL19 from accessing PhoQ in the periplasm, a barrier function that has been observed for other AMPs^48,49^. Alternatively, CCL19 may exhibit salt sensitivity, as has been observed for other AMPs^50,51^, or its fusion to the SLAY surface display protein may have altered its activity. However, given that the PhoQ-activating fragment of CCL19 is located at its C terminus (**Figure 4**), it is unlikely that context dependency at the CCL19 N terminus is responsible for the discrepancy in activation that we observed between our *E. coli* and *S*. Typhimurium experiments. Future studies could examine whether any of these effects, or other differences between *S*. Typhimurium and our engineered *E. coli* strain, explain the inability of CCL19 to activate PhoPQ in *S*. Typhimurium. These factors could also explain the different orders of activation strength that we observed between AMPs when surface-displayed in our *E. coli* strain and when supplied exogenously to *S*. Typhimurium.

Our studies of PhoPQ orthologs suggest that AMP specificity may have evolved across different species of pathogenic Enterobacteriaceae. While *S*. Typhimurium, ExPEC, and *K. pneumoniae* can all colonize the human gut and cause systemic infections, they can each infect distinct sets of tissues within the human body and interact differently with the human immune system^8,52–55^. For example, ExPEC commonly causes bloodstream and urinary tract infections (UTIs)^52^, whereas *Salmonella* species predominantly cause gastroenteritis and systemic infections^52,54^. Like ExPEC, *K. pneumoniae* can cause UTIs and bloodstream infections, but unlike ExPEC, *K. pneumoniae* is also commonly associated with pneumonia^53^. AMPs are also differentially expressed throughout the human body (**Supplementary Figure 6a**) and play varying roles in immunity that could affect their interactions with pathogenic bacteria. Through their unique infection lifestyles, pathogenic Enterobacteriaceae may regularly encounter different sets of AMPs, which could place evolutionary pressure on PhoQ to sense AMPs most relevant to their specific infection niches. Future studies will be required to validate activation of ExPEC and *K. pneumoniae* PhoPQ in their native contexts by human AMPs.

Finally, SLAY-TCS should be directly extensible to other peptide-sensing TCSs. Many AMP-sensing TCSs are associated with virulence, including CsrRS from Group A streptococci^56^, *Pseudomonas aeruginosa* ParRS, CprRS, and PmrAB^57–59^, *Vibrio cholerae* VprAB^60^, and *Staphylococcus aureus* ApsSR^61^. SLAY-TCS could be used to examine the responses of these TCSs to human AMPs, which could uncover new avenues for host-pathogen interaction in these organisms. This approach could also be used to examine TCSs involved in interbacterial interactions, such as bacteriocin-sensing TCSs of lactic acid bacteria involved quorum sensing and bacterial warfare^62^. By enabling high throughput characterization of peptide-TCS interactions, SLAY-TCS can shed new light on a wide range of microbiological pathways.

## Methods

### Bacterial strains and plasmids

Bacterial strains used in this work are listed in **Supplementary Table 5**. Genomic integration cassettes were introduced into *E. coli* BW30007 using clonetegration^63^ to produce strains KB1, KB1 PhoQ H277A, AM1, and AM2. Bacterial strains are available upon request.

Plasmids used in this work are listed in **Supplementary Table 6** and are available from Addgene (identifiers listed in **Supplementary Table 6**).

### Growth media

All *E. coli* experiments were conducted in M9 media containing 1x M9 salts, 0.4% glucose, and 0.2% casamino acids, adjusted to pH 7.3 and supplemented with 1 mM MgCl_2_, unless otherwise specified. All *S*. Typhimurium experiments were performed in N-minimal media^64,65^ containing 1x Buffer N (5 mM KCl, 7.5 mM (NH_4_)_2_SO_4_, 0.5 mM K_2_SO_4_, 1 mM KH_2_PO_4_, 100 mM Tris-HCl), 0.4% glucose, 0.1% casamino acids, and 2 mg/L thiamine hydrochloride, adjusted to pH 7.3 and supplemented with 1 mM MgCl_2_, unless otherwise specified. Kanamycin (50 μg/mL) and ampicillin (100 μg/mL) were added as necessary for maintenance of genomic inserts or plasmids.

### *E. coli* experiments

For all *E. coli* experiments except the sort-seq screen, 14 mL culture tubes containing 3 mL M9 media were inoculated from −80°C glycerol stocks and grown overnight at 37°C and 250 r.p.m. For each strain, 15 μL of overnight culture was used to inoculate 14 mL culture tubes containing 3 mL of fresh M9 media, which were grown at 37°C and 250 r.p.m for 2 h. The optical density (OD_600_) of these cultures was determined using a Cary50 ultraviolet-visible spectrophotometer (Agilent, Inc.). Cultures were subsequently diluted to OD_600_ = 0.0001 (peptide display and magnesium dose response experiments) or OD_600_ = 0.05 (LL-37 dose response experiments) in 200 μL cultures in 96-well plates. Isopropyl-β-D-1-thiogalactoside (IPTG), MgCl_2_, or LL-37 were supplemented as indicated, plates were sealed with adhesive aluminum foil covers, and cultures were grown at 37°C and 900 r.p.m. for 4.5 h (peptide display and magnesium dose response experiments) or for 1.25 h (LL-37 dose response experiments). Endpoint absorbance at 600 nm was measured using a Tecan Infinite M200 PRO plate reader, and measurements were converted to OD_600_ using a standard curve. Plates were placed on ice to arrest growth and cells were diluted into PBS for analysis by flow cytometry.

### *S*. Typhimurium experiments

14 mL culture tubes containing 3 mL N-minimal media were inoculated from −80°C glycerol stocks and grown overnight at 37°C and 250 r.p.m. For each strain, 50 μL of overnight culture was used to inoculate 14 mL culture tubes containing 3 mL of fresh N-minimal media, which were grown at 37°C and 250 r.p.m for 2 h. The optical density (OD_600_) of these cultures was determined using a Cary50 ultraviolet-visible spectrophotometer (Agilent, Inc.). Cultures were subsequently diluted to OD_600_ = 0.05 in 96-well plates at 50 μL (human AMP on/off experiments) or 200 μL (LL-37 and TCP C-terminal peptide dose response curves) culture volumes, with peptides supplemented as indicated (peptide commercial source information in **Supplementary Table 7**). Plates were sealed with adhesive aluminum foil covers and cultures were grown at 37°C and 250 r.p.m. for 1.25 h. Plates were placed on ice to arrest growth and cells were diluted into PBS for analysis by flow cytometry.

### Flow cytometry analysis

For all experiments except the sort-seq experiment, flow cytometry data acquisition was performed using a BD FACScan (Becton Dickinson) and FlowJo software (FlowJo, LLC). The flow cytometer was fitted with custom blue (488 nm, 30 mW) and yellow (561 nm, 50 mW) solid-state lasers (Cytek Biosciences) and with a custom 510/21 nm FL1 emission filter.

Side scatter (SSC) thresholds of 59% and 53% were used to separate cells from debris for *E. coli* and *S*. Typhimurium samples, respectively. 30,000 events were collected for each sample where possible; fewer events were collected for some samples with very low cell density. Fluorescent beads (Rainbow Calibration beads, Spherotech, RCP-30-5A) samples were also acquired during every flow cytometry experiment to enable conversion of data from arbitrary units (a.u.) to molecules of equivalent fluorescein (MEFL). For beads samples, 10,000 events were collected and an SSC threshold of 20% was used to separate beads from debris.

Flow cytometry data was analyzed using the FlowCal^66^ Python interface. For all experiments except the sort-seq screen, beads were gated using a 30% density gate in forward scatter (FSC) and SSC and used to generate a calibration curve to convert from a.u. to MEFL units. *E. coli* samples were gated using an 85% density gate in FSC and SSC. *S*. Typhimurium samples were first gated to remove samples with very low FSC (FSC < 1.5 a. u.) and then gated using an 85% density gate in FSC and SSC. Fluorescence values were converted to MEFL using the beads calibration curve (**Supplementary Figure 9a**).

Each reported fluorescence value represents the arithmetic mean of all gated events in a given sample. Fold change was calculated using autofluorescence-subtracted mean fluorescence values, in which the mean fluorescence of an autofluorescence control was subtracted from the mean fluorescence of the sample to isolate the fluorescent protein-derived signal. Fold change values were calculated by dividing the autofluorescence-subtracted mean fluorescence of a sample by the autofluorescence-subtracted mean fluorescence of an appropriate negative control. Best-fit Hill curves and associated half-maximal activation values were calculated using the lmfit Python package^67^. The reported error in half-maximal activation is the standard error as calculated by lmfit^67^.

### Human AMP library design and construction

#### Oligo pool design

First, the Antimicrobial Peptide Database (APD3)^17^ was queried using *Homo sapiens* in the species field to generate a list of all human AMPs. At the time of the query, there were 133 AMPs that met this criterion (seven more AMPs have been added to the database since we conducted the original query). 117 of these were within 104 amino acids in length, the maximum length compatible with our library construction approach, which was limited by restrictions on the length of commercial oligos. We generated oligo sequences corresponding to each of these 117 human AMPs, along with 5 no peptide control sequences in which a stop codon is used in place of a peptide (**Supplementary Table 1**). Five control sequences were included to ensure adequate library representation, since the no peptide control forms the basis for all fold change calculations in our sort-seq experiment. All oligo sequences were designed to be the same length (350 bp) to reduce amplification bias during polymerase chain reaction (PCR) steps involved in library construction and in amplicon generation for Illumina next-generation sequencing (NGS). For most AMPs, a non-coding DNA sequence was included after the stop codon to adjust for differences in AMP length while maintaining constant oligo length.

#### Oligo pool amplification

We ordered the human AMP oligos from Integrated DNA Technologies (Integrated DNA Technologies) as an oPool. oPools contain a substantial proportion of short oligo fragments; thus, we used PCR to select for fulllength oligos. Oligos in the oPool were originally designed to contain only one primer binding site on their 3’ end. To enable PCR amplification, we introduced a second primer binding site on the 5’ end of the oligos using a ligation reaction. We designed custom DNA adapters consisting of two partially complementary oligos to facilitate this ligation reaction (**Supplementary Figure 10**). One “extension” oligo consisted of the sequence to be ligated to the 5’ end of the oligos in the oPool (KB636: 5’-GGTACTGTTGATGTAGGTTCACGATCCT-3’). A second “bridge” oligo contained regions complementary to both the first oligo and to a short, common sequence designed into the 5’ end of all oligos in the oPool (KB637: 5’-TGTTCGGAGACCAAGGATCGTGAACCTACATCAACAGTACC-3’). This bridge oligo promotes adapter binding to oPool oligos with intact 5’ ends, further enhancing selection for full-length oligos. First, oPool oligos were phosphorylated at their 5’ ends using T4 Polynucleotide Kinase (New England Biolabs, Ipswich, MA). DNA adapters were then annealed to the phosphorylated oPool oligos by incubating extension, bridge, and oPool oligos at 95°C and decreasing the incubation temperature to 55°C at a rate of 1°C/min. The annealed oligos were incubated with T4 DNA ligase (New England Biolabs) to covalently link the extension oligos to the oPool oligos. Phusion PCR reactions (New England Biolabs) were used to amplify these extended oligos, with forward and reverse primers introducing BsaI cut sites for use in a subsequent Golden Gate assembly reaction (KB614: 5’-TGGTCTCTGCTGTATCATCTGCGTACTCCATTA-3’ and KB631: 5’-GTTCACGATCCTTGGTCTCCGAACA-3’).

#### Backbone amplification

Phusion PCR reactions (New England Biolabs) were used to generate backbone fragments for library assembly. A previously-generated peptide display plasmid (pKB225) was used as a template, and forward and reverse primers were used to add BsaI cut sites (KB554: 5’-AGGTCTCAGTTCCTCCGATACCCGCAG-3’ and KB626: 5’-AGGTCTCACAGCTGCCTGGCGGCAGTAGCGC-3’). Backbone PCR amplicons were treated with BsaI and DpnI (New England Biolabs) to generate overhangs for oligo insertion and to remove any template plasmid, respectively. Amplicons were subsequently treated with calf intestinal phosphatase (Quick CIP, New England Biolabs) to remove 5’- and 3’-phosphate groups to prevent backbone self-ligation.

#### Plasmid library assembly and purification

A two-part Golden Gate assembly reaction with BsaI (New England Biolabs) was used to insert the PCR-amplified oligo pool into the PCR-amplified backbone. This assembly was transformed into NEB 10-β competent cells via electroporation. As a control to estimate background, we also transformed a mock assembly reaction in which the insert was omitted. We estimated our background to be 0.5% based on the transformation efficiencies of our assembly and control reactions (313.6 cfu/ng and 1.4 cfu/ng, respectively). We scraped approximately 27,000 colonies (221x library coverage) from our assembly reaction transformation into LB and immediately purified their plasmid DNA using the Plasmid Plus Midi Kit (Qiagen) according to the manufacturer’s instructions.

#### Transformation into KB1

We transformed the purified library plasmid DNA into KB1 via electroporation and scraped approximately 28,400 colonies (233x library coverage) resulting from this transformation into 50 mL M9 media (not supplemented with MgCl_2_). Cells were pelleted at 4000 r.p.m. and 4°C for 10 min and the supernatant was decanted. Cells were similarly resuspended in 25 mL M9 media (not supplemented with MgCl_2_) and pelleted twice before being finally resuspended in 25 mL M9 media (not supplemented with MgCl_2_). Approximately 4 mL of cells were dispensed into 50 μL aliquots that were immediately frozen at −80°C for use in inoculating future experimental cultures. The remaining cells (~21 mL) were pelleted (4000 r.p.m. and 4°C for 10 min), the supernatant was decanted, and the pellet was frozen at −30°C. DNA purification was later performed for these pelleted cells using the Plasmid Plus Midi Kit (Qiagen) according to the manufacturer’s instructions. This purified DNA was used as a reference (pre-sort sample) to determine whether any growth bias occurred during the sort-seq experiment.

### Sort-seq experiment

#### Growth

Autofluorescence, no peptide, and C18G control strain cultures were inoculated from −80°C glycerol stocks into 14 mL culture tubes containing 3 mL M9 media. After growth overnight at 37°C and 250 r.p.m., 15 μL of each overnight culture was used to inoculate a 14 mL culture tube containing 3 mL of fresh M9 media, which were grown at 37°C and 250 r.p.m for an additional 2 h. For the human AMP library, one 50 μL frozen aliquot was used to inoculate a 14 mL culture tube containing 3 mL M9 media, which was grown at 37°C and 250 r.p.m for 2 h alongside the control strain cultures. The optical density (OD_600_) of these cultures was determined using a Cary50 ultraviolet-visible spectrophotometer (Agilent, Inc.). Each control strain and the human AMP library were diluted to OD_600_ = 0.001 in 14 mL culture tubes containing 3 mL fresh M9 media supplemented with 500 μM isopropyl-β-D-1-thiogalactoside (IPTG). Cultures were grown at 37°C and 250 r.p.m for 3 h and then placed on ice to arrest growth.

#### Sorting and DNA purification

Flow cytometry measurements and cell sorting were performed using a Sony SH800S cell sorter. A forward scatter (FSC) threshold of 0.1% was used to separate cells from debris. The human AMP library was gated using hand-drawn FSC and back scatter (BSC) gates to separate cells from any additional debris (retained approximately 99.6% of events) and then sorted into eight fluorescence bins, spaced approximately logarithmically (**Supplementary Figure 9b**, **Supplementary Table 2**). 100,000 cells were collected for each fluorescence bin, along with a control bin in which no fluorescence gating was applied. Cells were sorted into 3 mL cold LB media, which was then added to 47 mL room-temperature LB for a total culture volume of 50 mL. Each culture was grown for approximately 6 hours at 37°C and 250 r.p.m. until reaching an OD_600_ = 0.3 – 0.4. Cells were pelleted and pellets were stored at −30°C overnight. Minipreps were performed to isolate plasmid DNA from the cell pellets, and DNA concentrations were measured using a Qubit 4 Fluorometer (Invitrogen).

#### Sort-seq flow cytometry analysis

Flow cytometry measurements were analyzed using FlowCal^66^ to generate the histograms in **Figure 3b** and **Supplementary Figure 5a**. No gating was applied to these samples during collection or analysis. The human AMP library data in **Figure 3b** is based on two measurements of the same sample consisting of 100,000 events each, for a total of 200,000 events. 30,000 events were collected for all samples shown in **Supplementary Figure 5a**. Fluorescence values are reported in arbitrary units (a.u.).

### Next-generation sequencing

Phusion PCR reactions (New England Biolabs) were used to generate amplicons for Illumina NGS from purified plasmid DNA. For each sample, two 50 μL reactions were performed using 100 ng of plasmid DNA template and 0.2 μM each of forward and reverse primers (Integrated DNA Technologies) (**Supplementary Table 8**). For the pre-sort sample, forward and reverse primer overhangs introduced partial Illumina adapters to enable compatibility with the GENEWIZ Amplicon-EZ service. For post-sort samples, forward and reverse primers were used to introduce unique 10 bp barcode sites to enable multiplexed sequencing and bin assignment during NGS data processing. After an initial denaturation step of 30 s at 98°C, 12 amplification cycles were performed of 10 s at 98°C, 30 s at 68°C, and 15 s at 72°C. A final elongation step was performed for 10 min at 72°C.

For each sample, the two PCR reactions were combined and subsequently purified using spin columns (EconoSpin). DNA concentrations were measured using a Qubit 4 Fluorometer (Invitrogen) and concentrations were adjusted for Illumina NGS sequencing. The pre-sort sample was sequenced using the GENEWIZ Amplicon-EZ service. Sort-seq samples (Bin 1-8 and no fluorescence gate samples) were combined in equimolar amounts to produce a single sample, which was sent to GENEWIZ for library preparation and sequencing.

Subsequent steps were performed at the GENEWIZ facility through their commercial services. DNA library preparation, clustering, and sequencing were performed using NEBNext Ultra DNA Library Prep kit following the manufacturer’s recommendations (Illumina). For the pre-sort sample, adapter-ligated DNA was indexed and enriched by limited cycle PCR. For the sort-seq sample, the DNA library was validated using a TapeStation (Agilent Technologies) and quantified using a Qubit 2.0 Fluorometer (Invitrogen) and qPCR. For both pre-sort and sort-seq samples, DNA was loaded on the Illumina MiSeq instrument according to the manufacturer’s instructions and sequenced using a 2×250 paired-end configuration. Image analysis and base calling were conducted by the MiSeq Control Software on the Illumina MiSeq instrument. Raw Illumina reads were checked for adapters and quality via FastQC and trimmed of their adapters using Trimmomatic v0.36. Raw sequence data (.bcl files) was converted to fastq files and de-multiplexed using Illumina bcl2fastq.

### Sort-seq data analysis

Paired-end reads were combined using FLASH 1.2.11^68^. Each paired-end read was assigned to a fluorescence bin based on the barcode introduced during PCR amplification using custom code and the Python regex package^69^. The peptide amino acid sequence encoded in each paired-end read was determined using BioPython^70^. We used the frequency of reads encoding each peptide sequence in each bin and the distribution of events collected during sorting (**Supplementary Table 2**) to estimate the mean fluorescence for each peptide. We assumed a lognormal distribution, consistent with previous literature^71^. To eliminate low-quality data, we excluded peptide sequences from further analysis based on three criteria: fewer than 100 paired-end reads across all eight fluorescence bins, low confidence in our mean fluorescence calculation based on bootstrapping analysis, and lower fluorescence fold change than an autofluorescence-only control. For additional details, see **Supplementary Note 1**.

### Human AMP clustering

Agglomerative clustering was performed using USEARCH^72^ to group related human AMPs. Clustering was performed using average linkage, which joins clusters based on the average pairwise distance between peptides in each cluster. A 70% distance threshold was found empirically to yield reasonable AMP clusters.

### Calculation of peptide biochemical and biophysical descriptors

We calculated 1,284 biochemical and biophysical descriptors for all peptides in our library that met our quality control criteria and were greater than 10 amino acids in length, as summarized in **Supplementary Table 3**. Basic features were calculated using custom code, BioPython^70^, or built-in Python functions. Pairwise relative amino acid composition was calculated using custom code as in Lee *et al*. 2016^73^. The remaining descriptors were calculated using propy^74^. For additional details, see **Supplementary Note 2**.

### Development and assessment of a cathelicidin sparse robust linear model

The longest contiguous subsequence algorithm was used to assign each peptide variant in the human AMP library to a parent human AMP. Using this approach, 410 peptides in our dataset were found to be cathelicidin-like. Of these cathelicidin-like peptides, 25% (102 peptides) were withheld as a test set for model validation, with the remaining 308 peptides constituting a training set. The previously calculated biochemical and biophysical descriptors served as inputs to the model and the logarithm of the fold change of activation from the sort-seq experiment served as the target variable. The training set was used for feature selection and to train a sparse robust linear model^43^, with fourteen features ultimately selected as shown in **Supplementary Table 4**. Model performance was evaluated using the Pearson correlation coefficient (*r*) and coefficient of determination (*r*^2^) between the predicted log_10_(Fold Change) values from the model and the measured log_10_(Fold Change) values from the sort-seq experiment. These coefficients were calculated using scipy^75^ and Scikit-learn^76^, respectively. For additional details, see **Supplementary Note 3**.

### Protein sequence alignments and identification of transmembrane domains for PhoQ orthologs

Protein sequence alignments of PhoQ orthologs were conducted using MEGA-X^77^. The ClustalW algorithm was used with a gap opening penalty of 10 and gap extension penalty of 0.2. Percent identity was calculated based on these alignments. Transmembrane domains were identified using TMHMM v2.0^78^.

### Peptide structural data

Peptide structures were downloaded from the RCSB Protein Data Bank^79^. Image renderings were performed using UCSF Chimera^80^.

### Human AMP tissue expression data

Tissue-based RNA expression data was downloaded from the Human Protein Atlas RNA consensus tissue gene data dataset^33,41^. Human Protein Atlas data is available from http://v20.proteinatlas.org. Peripheral blood mononuclear cell (PBMC) single-cell RNA expression data^42^ was downloaded from the Gene Expression Omnibus database under accession numbers GSM3454528 (naïve cells) and GSM3454529 (*Salmonella*-exposed cells). RNA transcript counts were normalized to the total number of sequencing reads for a given cell as indicated by the cell barcode.

### Western blot analysis of AMP expression in THP-1 and human monocyte-derived macrophages (hMDMs)

Preparation of hMDMs and THP-1 cells was described previously^81,82^. hMDMs and THP-1 infection with *S*. Typhimurium and Western blot analysis were performed as previously described^83^. Macrophage cell lysates were collected at 4 h post-infection and subjected to Western blot for GNLY (R&D Systems Cat# AF3138-SP), CCL20 (R&D Systems Cat# AF360-SP), and/or actin (Abcam Cat# 179467) using 40 μg of total protein.

## Data availability

The raw and processed NGS datasets that support the findings of this study (as reported in **Figure 3c-f** and **Supplementary Figure 5b-c,f**) have been deposited in the Gene Expression Omnibus (GEO) Database with the series accession ID GSE174191 (https://www.ncbi.nlm.nih.gov/geo/query/acc.cgi?acc=GSE174191). Raw flow cytometry data for the following figures is available from the corresponding author upon reasonable request: **Figure 1c-d, Figure 2d, Figure 3b, Figure 4b, Figure 5a-b,d, Figure 6b, Supplementary Figure 2, Supplementary Figure 5a,d**, and **Supplementary Figure 8b**.

Raw OD_600_ data for the following figure is available from the corresponding author upon reasonable request: **Supplementary Figure 3**.

Raw Western blot data for the following figures is available from the corresponding author upon reasonable request: **Supplementary Figure 6c-d**.

Human AMP sequences were downloaded from the Antimicrobial Peptide Database (APD3)^17^. Peptide structures were downloaded from the RCSB Protein Data Bank^79^.

Tissue-based RNA expression data was downloaded from the Human Protein Atlas RNA consensus tissue gene data dataset^33,41^. Human Protein Atlas data is available from http://v20.proteinatlas.org.

Peripheral blood mononuclear cell (PBMC) single-cell RNA expression data^42^ was downloaded from the Gene Expression Omnibus database under accession numbers GSM3454528 (naïve cells) and GSM3454529 (*Salmonella*-exposed cells).

## Code availability

Code for the analysis and visualization of sort-seq data, except for the cathelicidin machine learning model, is available on GitHub at https://github.com/krbrink/PhoPQ_hAMP_sort-seq. Code for the cathelicidin machine learning model is available on GitHub at https://github.com/kennygrosz/PhoPQ_Activation_model.

## Ethics statement

This study was carried out in accordance with the recommendations of Nationwide Children’s Institutional Review Board with written informed consent from all subjects. The protocol was approved by Nationwide Children’s Institutional Review Board.

## Acknowledgements

We thank Bryan Davies for generously providing SLAY source plasmids and insights into using SLAY in *E. coli*. The following reagent was obtained through BEI Resources, NIAID, NIH: *Salmonella enterica* subsp. *enterica*, Strain 14028s Δ*phoQ* (Serovar Typhimurium), NR-40554. We would also like to thank Joel Moake for the use of his cytometer. This work was supported by the Welch Foundation (C-1856), NSF CAREER (1553317), and NIH/NIAID 1R01AI155586-01A1. K.R.B. was supported by the Nettie S. Autrey Fellowship from Rice University. K.V.H. and J.G. were supported by funds from Nationwide Children’s Hospital.

## Author contributions

K.R.B. and J.J.T. conceived of the project. J.J.T. and J.S.G. supervised the project. K.R.B. and A.M.M. (identification of PhoPQ reporter promoter, PhoPQ orthologs), K.V.H. (AMP expression in human cell lines), and K.R.B. (all other experiments) designed and executed experiments. K.G. and K.R.B. designed, implemented, and analyzed the cathelicidin machine learning model. K.R.B. analyzed reported results and generated figures and tables. K.R.B. and J.J.T. wrote the manuscript.

