## Supplementary Information for "High-throughput discovery of peptide activators of a bacterial sensor kinase"

**Supplementary Note 1.** Sort-seq data analysis.

*Estimation of mean fluorescence output from FACS and NGS data*

All data analysis was performed using custom code in Jupyter notebooks running Python 3.7.4. Forward and reverse reads were combined using FLASH 1.2.11[1] using a maximum overlap of 150 bp, minimum overlap of 10 bp and maximum mismatch density of 0.1. Each forward or reverse read was assigned to a bin based on its best match to the primer barcodes introduced during PCR amplification, as determined using regular expressions via the regex package[2]. A bin assignment was considered valid if a read sequence and its best-match barcode differed by 2 or fewer errors. Any paired-end read where the best-match forward and reverse barcodes corresponded to different bins was excluded from further analysis. Similarly, any paired-end read where neither the forward nor reverse reads could be assigned to a bin was excluded. Paired-end reads where the forward read contained a valid barcode sequence but the reverse read did not, and vice versa, were assigned the bin of the valid barcode.

For paired-end reads with valid bin assignments, the peptide-encoding DNA sequence was identified based on its proximity to a sequence encoding the reverse primer binding site. Only paired-end reads that contained exactly one reverse primer binding site (with up to 2 errors) were considered valid and retained for analysis. For each valid paired-end read, the peptide-encoding DNA sequence was translated *in silico* using BioPython[3]. Reads that encoded a peptide sequence but did not include a stop codon were excluded from downstream analysis. All valid paired-end reads corresponding to the same peptide translation and bin assignment were cross-tabulated. This included reads encoding no peptide controls, which were analyzed collectively as a single no peptide control. These counts were normalized by the total number of paired-end reads in each bin to obtain frequency values. Frequency values were weighted based on the relative fraction of flow cytometry events that occurred in each fluorescence bin (**Supplementary Table 2**) to estimate the original fluorescence distributions. These distributions were transformed into probability-normed space for plotting by dividing by the width of each fluorescence bin.

We used the empirical distributions described above to estimate the mean fluorescence output for each displayed peptide in the library. Consistent with previous literature, we assumed that fluorescence distributions were lognormal in shape[4]. First, we calculated the lognormal distribution parameters $\hat{\mu}$ and $\hat{\sigma}^2$. We denote the probability of an event falling in bin *j* as $r_j$ and specify the total number of fluorescence bins as *m* such that $\sum_{j=1}^{m} r_j = 1$. We identified the center of each fluorescence bin *j* in logarithmic space, given by $\beta_j$, from its lower and upper boundaries $L_j$ and $U_j$, respectively).

$$\beta_j = \frac{\ln(L_j) + \ln(U_j)}{2}$$

We calculated $\hat{\mu}$ and $\hat{\sigma}^2$ assuming that all events in a given bin *j* exhibited a fluorescence $\beta_j$.

$$\hat{\mu} = \sum_{j=1}^{m} r_j \beta_j$$

$$\hat{\sigma}^2 = \sum_{j=1}^{m} r_j (\beta_j - \hat{\mu})^2$$

Finally, we used the parameters $\hat{\mu}$ and $\hat{\sigma}^2$ to calculate mean fluorescence $\hat{v}$.

$$\hat{v} = \exp\left(\hat{\mu} + \frac{\hat{\sigma}^2}{2}\right)$$

We found that this approach performed comparably to a maximum likelihood estimator (MLE) approach[4] and required less computation time. Fold change values were determined by dividing the mean fluorescence for a given peptide by the mean fluorescence of the no peptide control.

*Quality control*

We performed three steps to exclude peptides with low-quality data from our analysis. First, we removed peptides with fewer than 100 paired-end reads in total across all eight fluorescence bins. Next, we performed bootstrapping to assess the robustness of our calculated fluorescence means. High variability across different bootstrapping iterations indicates low confidence in the result for a given peptide. We performed 1,000 bootstrapping iterations in which we recreated our sort-seq results *in silico*. In each iteration, we populated each fluorescence bin with peptides chosen randomly with replacement based on their frequency in the original dataset. We repeated our calculation of mean fluorescence as described above for each simulated dataset and examined the distribution of mean fluorescence values across all bootstrapping iterations for each peptide. We summarized the variability across iterations based on the width of the $5^{th}$ to $95^{th}$ percentile range of mean fluorescence values. We divided this range by the mean fluorescence value calculated for the original dataset to obtain a normalized $5^{th}$-$95^{th}$ percentile range. Peptides with a normalized $5^{th}$-$95^{th}$ percentile range greater than or equal to 0.4 were excluded from subsequent analyses. Finally, we excluded any peptides with a fold change less than or equal to the fold change that we observed for an autofluorescence-only control that was measured on the same day as the sort-seq screen. This last criterion eliminates peptides that likely have mutated copies of the fluorescent mNeonGreen reporter.

**Supplementary Note 2.** Calculation of peptide biochemical and biophysical descriptors.

Charge was calculated by assigning each lysine (K) and arginine (R) residue a charge of +1 and each aspartic acid (D) and glutamic acid (E) residue a charge of -1, with all other residues assigned a charge of 0. Length was calculated using the built-in Python len function. Grand average of hydropathicity (GRAVY), isoelectric point, molecular weight, aromaticity, and instability were calculated using built-in functions in BioPython[3].

In addition to the seven descriptors above, 1,277 additional descriptors were calculated to generate and assess the performance of the cathelicidin sparse robust linear model, for a total of 1,284 descriptors (**Supplementary Table 3**). These descriptors can be classified into four feature types: residue composition, autocorrelation, physicochemical composition, and sequence order features. Autocorrelation and sequence order features rely on lag or distance ($\delta_{max}$ or $\lambda_{max}$) parameters that can only be calculated for peptides with length $L > \delta_{max}$ and $L > \lambda_{max}$. To enable the use of these features, we limited our dataset to peptides greater than 10 amino acids in length and set $\delta_{max} = 10$ and $\lambda_{max} = 10$ for all autocorrelation and sequence order features.

Residue composition features included amino acid composition, dipeptide composition, and pairwise relative amino acid composition. Amino acid and dipeptide compositions measure the frequencies of amino acid residues and dipeptides (two contiguous residues) across a peptide sequence. These features were calculated using propy[5]. Pairwise relative amino acid composition measures the occurrences of a given amino acid relative to those of a second amino acid for all pairwise combinations of amino acids. Pairwise relative amino acid composition was calculated as in Lee *et al.* 2016[6].

Autocorrelation features measure the similarity in properties between residue pairs separated by a distance δ in the peptide primary sequence. All autocorrelation features were calculated using propy[5]. Following Ong *et al.* 2007[7], we included eight descriptors in our autocorrelation calculations: average flexibility, free energy, hydrophobicity, mutability, polarizability, residue accessible surface area (ASA), residue volume, and steric parameters. Each of the three autocorrelation features included in this analysis differs based on its approach to calculating similarity. Normalized Moreau-Broto autocorrelation uses the product of property values of residues separated by a distance δ, Moran autocorrelation calculates the product of the differences between those property values and the average across the peptide sequence, and Geary relies on the squared difference between property values for residues separated by a distance δ[7].

Physicochemical composition features were calculated using propy[5] and include composition, transition, and distribution (CTD) for each of seven descriptors: charge, hydrophobicity, normalized van der Waals volume (VDWV), polarity, polarizability, secondary structure, and solvent accessibility. For each descriptor, the 20 amino acids are divided into three groups based on their physicochemical properties[5]. Composition (C) measures the frequency of residues of a given group in a peptide sequence. Transition (T) measures the frequency of contiguous residue pairs where adjacent residues belong to different groups. Distribution (D) measures the fractional location at which the first, 25$^{th}$ percentile, 50$^{th}$ percentile, 75$^{th}$ percentile, and 100$^{th}$ percentile residues in a given group occur across the length of the peptide.

Similar to autocorrelation features, sequence order features consider the ordering of residues in the primary sequence of a peptide. Sequence-order-coupling number uses a distance matrix to assess physicochemical distances between different amino acid residues separated by a primary sequence distance δ. We measured sequence-order-coupling number using both Grantham[8] and Schneider-Wrede[9] distance matrices in propy[5]. Pseudo-amino acid composition (PAAC) and amphiphilic pseudo-amino acid composition (APAAC) were calculated according to Chou[10–12] using propy[5] with $\lambda_{max} = 10$. PAAC incorporates correlations between a parameter that contains hydrophobicity, hydrophilicity, and side-chain mass components. APAAC considers correlations in hydrophobicity and hydrophilicity. We used the propy default weights of 0.05 for PAAC and 0.5 for APAAC.

**Supplementary Note 3.** Development and assessment of a cathelicidin sparse robust linear model.

*Identifying parent human AMPs of peptide variants in the human AMP library*

Each peptide sequence longer than 10 amino acids in length (3,495 sequences) was compared pairwise with each of the 117 sequences designed into the human AMP library using the longest contiguous subsequence (LCS) algorithm, which was sourced from the Subsequences package in Julia[13]. This algorithm returns the longest common shared sequence between two sequences.

We considered a peptide variant to be derived from a given parent human AMP sequence if their longest common contiguous subsequence exceeded an LCS threshold *t*. The peptide was then considered to belong to the same cluster as the parent human AMP. We identified the smallest value of *t* for which no peptide matched human AMPs from two or more human AMP clusters. This value was found to be $t = 7$.

At this level of *t*, 748 peptides were not able to be matched to any parent human AMP cluster and were discarded from the dataset. The cathelicidin-like cluster was used to create an activation prediction model, and all other clusters were used to evaluate the generalizability of the resulting predictions.

*Feature selection*

First, 25% of the 410 cathelicidin-like peptides were randomly held out from the dataset for use in testing the performance of the model. The remaining 308 peptides (training set) were used for feature selection and model generation as described below.

A feature selection process was undertaken to maintain the relevant features while removing redundant or irrelevant features for predicting activation of PhoPQ by cathelicidin-like peptides. First, 94 features with no variance were removed from the dataset. The remaining 1,190 features were checked for redundancy with other features in the dataset. 212 features with a Pearson correlation coefficient of $|r| \geq 0.95$ with another feature in the dataset were removed. For each pair of redundant features, the feature eliminated from the dataset was the one possessing the lower magnitude correlation with the response variable, $\log_{10}$(Fold change). The remaining dataset contained 308 cathelicidin-like peptides and 978 features.

*Predictive model*

A sparse robust linear model[14] was produced to predict PhoPQ activation by cathelicidin-like peptides. Mathematically, this model solves the following formulation:

$$\min_{w,b} \sum_{i=1}^{n} \|y_i - (w^T x_i + b)\|_2^2 + \frac{1}{2\gamma}\|w\|_2^2 \quad s.t. \ \ \|w\|_0 \leq k$$

The first term is the standard mean-squared error loss between the model's prediction, $w^T x_i + b$, and the actual $\log_{10}$(Fold change) of PhoPQ activation, $y_i$. The second term is a regularization term to increase robustness and prevent overfitting. The constraint explicitly restricts the selection of more than k features to have nonzero coefficients in the final model. This model can be seen as doing feature selection on the fly, only keeping a feature in the model if it is highly relevant to predict the response variable. Sparse linear regression models have been shown to be quite effective in recovering the true support in high-noise environments[15].

The model was trained using the Interpretable AI implementation of sparse and robust linear regression, Optimal Feature Selection Learner[16], and solved using the binary relaxation method[15], which is shown to have comparable results to the exact formulation but is more computationally tractable. The model was fully developed in the Julia programming language.

This model has two relevant hyperparameters: k, which controls the sparsity of the model, and $\gamma$, which controls regularization, wherein a lower value of $\gamma$ implies a higher degree of regularization. The optimal value for these parameters was determined as the value that maximizes the average validation accuracy ($r^2$) over 15 rounds of shuffled cross-validation, in which the model was trained on 75% of the training data and its accuracy is measured on the remaining 25%. The implementation of shuffled cross-validation used was from Scikit-learn[17].

Two phases of cross-validation were performed to thoroughly search the 2D hyperparameter space. The first was a coarse search over a wide range of the hyperparameters. In this phase, it was observed that ɣ = 0.005 yields models with higher mean validation $r^2$ and lower variance in the validation $r^2$ for almost all values of k. A fine-grained search was then performed which examined all integer values of k between 5 and 70 and five ɣ values between 0.0025 and 0.0075. These models each exhibited similar performance, with a standard deviation in validation $r^2 \approx 0.1$. Thus, we chose to set the hyperparameters to k = 14 and ɣ = 0.0075, which produce a model that balances interpretability and performance. This model has an average validation $r^2$ = 0.71 (93% of maximum observed average validation $r^2$) and standard deviation of validation $r^2$ = 0.08.

With the optimal hyperparameters chosen, the model was re-trained using the whole training set of 308 cathelicidin-like peptides. The 14 features ultimately selected by the model, along with the coefficients and feature importances given to those features, are seen in **Supplementary Table 4**. Feature importance is calculated based on the coefficients of the feature in the model, after normalization based on the magnitudes of different features, and then scaled to sum to 1[16].

Pearson correlation ($r$) and coefficient of determination ($r^2$) between $\log_{10}$(Fold Change) values measured in the sort-seq screen and $\log_{10}$(Fold Change) values predicted by the cathelicidin sparse robust linear model were calculated using scipy[18] and Scikit-learn[17] packages, respectively.

### Supplementary Figures

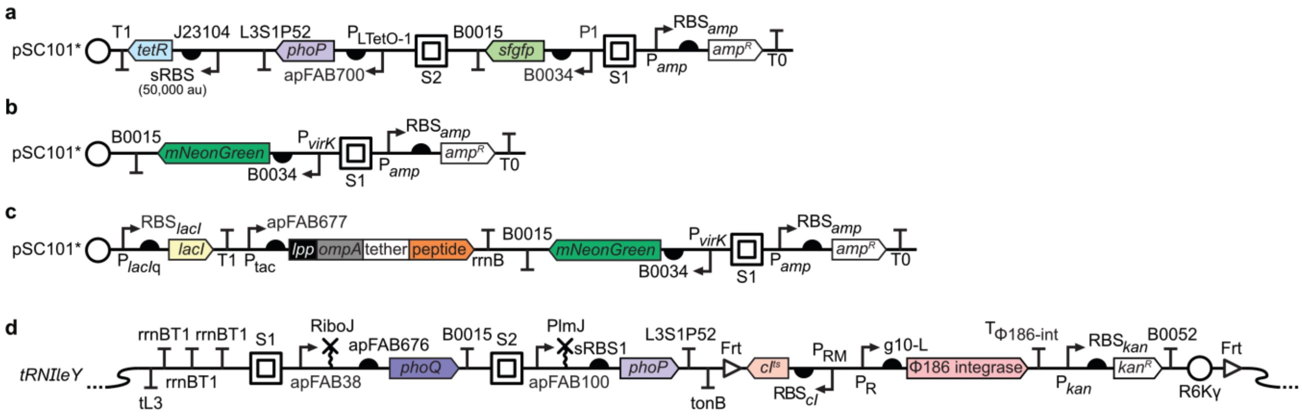


**Supplementary Figure 1.** Detailed diagrams of engineered genetic constructs.

(**a**) PhoP overexpression plasmid with *sfgfp* reporter. For reporter promoter identification experiment, P1 is the indicated PhoP-activatable promoter (*slyB*, *virK*, *ugtL*, *mgtC*, or *pmrD* promoter). P1 and $P_{LTetO-1}$ are insulated with inert spacer sequences S1 and S2 to reduce context dependency in promoter activity. (**b**) Diagram of $P_{virK}$-*mNG* reporter plasmid. (**c**) Diagram of peptide display plasmid, including $P_{virK}$-*mNG* reporter. (**d**) PhoPQ genomic expression cassette. Strains KB1, AM1, and AM2 encode *S.* Typhimurium, ExPEC EC958 (ST131), and *K. pneumoniae* Kp52145 *phoP* and *phoQ*, respectively. KB1 PhoQ H277A is identical to KB1 except that the histidine phosphorylation site of PhoQ has been mutated to an alanine residue. *phoP* and *phoQ* are each insulated with an inert spacer sequence (S1 and S2, respectively) and with a self-cleaving RNA element (RiboJ and PlmJ, respectively) to mitigate context-dependent effects on gene expression. Two different synthetic RBSs were used preceding *phoP* (sRBS1), with KB1 and AM2 using one synthetic RBS and AM1 using a second synthetic RBS. Additional gene expression cassettes ($cI^{ts}$ and Φ186 integrase) were required for genomic integration using clonetegration[19].

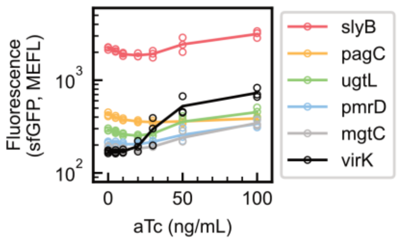


**Supplementary Figure 2.** Identification of a PhoP-activatable promoter to use as a reporter for PhoPQ activity.

Superfolder GFP (sfGFP) output from five candidate promoters was measured using flow cytometry in response to PhoP overexpression from an aTc-inducible promoter (**Supplementary Figure 1a**). Data was collected over $n$ = 3 separate days, with results from each day shown as a separate marker.

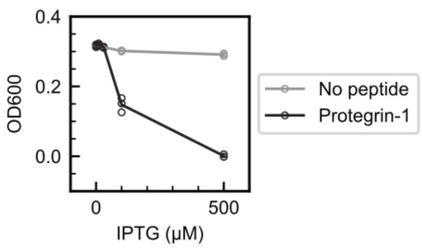


**Supplementary Figure 3.** Growth of KB1 PhoQ H277A strain displaying protegrin-1.

Endpoint OD600 measurements for the KB1 PhoQ H277A strain in response to IPTG induction of surface-displayed protegrin-1 and a no-peptide negative control. Measurements were performed after 4.5 h growth from an initial $OD_{600}$ = 0.0001. Data was collected over $n$ = 3 separate days, with results from each day shown as a separate marker.

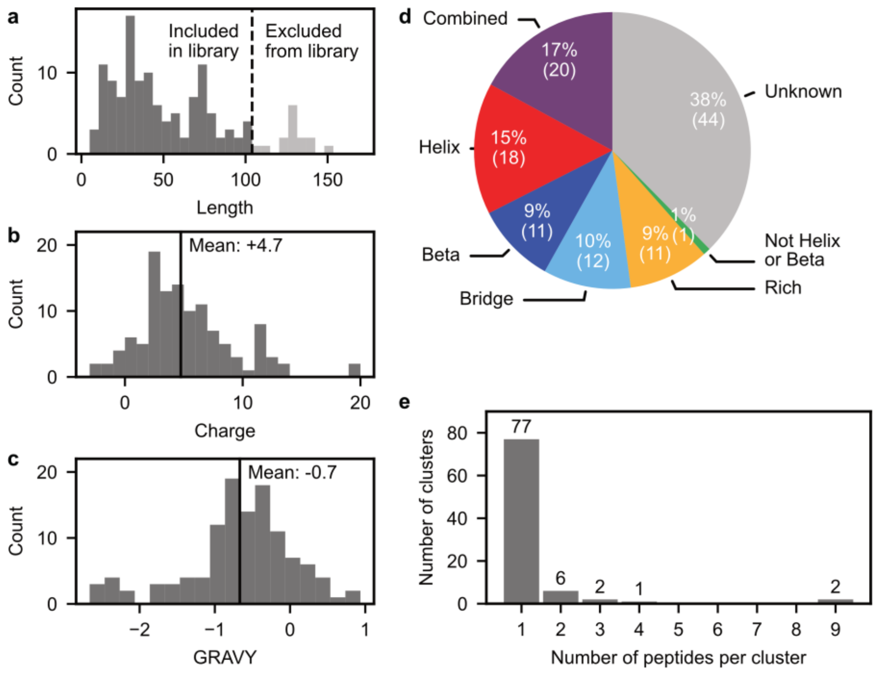


**Supplementary Figure 4.** Composition of the human AMP library.

(**a**) Lengths of all 133 human AMPs identified in APD3[20]. Peptides excluded from the human AMP library due to length considerations are indicated in light grey. (**b-d**) Charge, grand average of hydropathicity (GRAVY), and secondary structure distributions of AMPs in the human AMP library. (**e**) Distribution of cluster sizes for AMPs in the human AMP library.

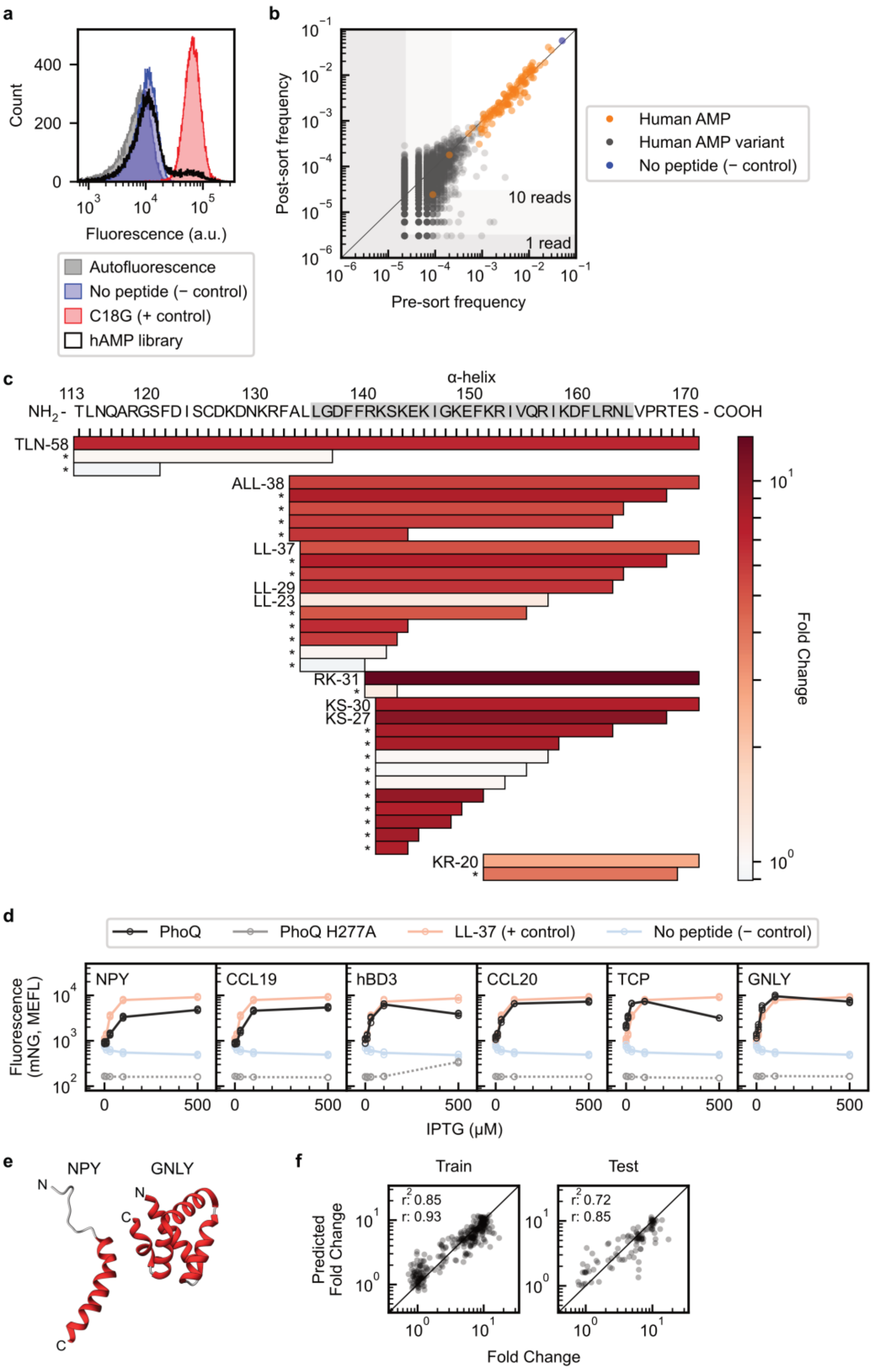


**Supplementary Figure 5.** Analysis and validation of sort-seq results for the human AMP library.

(**a**) mNeonGreen reporter expression in KB1 human AMP library (hAMP) prior to sorting. Overlaid histograms show fluorescence distributions for positive (C18G) and negative (no peptide) control peptide display strains, as well as an autofluorescence only control strain. Control strains were grown in single-strain cultures alongside the human AMP library. Each histogram represents the results of a single biological replicate ($n$ = 1). (**b**) Growth bias introduced in the sort-seq screen, as measured by peptide frequency in the human AMP library before and after the sort-seq workflow. Human AMPs are indicated in orange and variants in grey. No peptide control shown in blue. Pre- and post-sort frequency data was determined from single biological replicates (*n* = 1). (**c**) PhoPQ activation by human cathelicidin fragments of at least 3 amino acids in length in the sort-seq screen. Fragments are aligned to the TLN-58 sequence (* = human AMP variant). The α-helical region of LL-37 is indicated by the grey boxed region in the TLN-58 sequence. Data represents results from a single biological replicate (*n* = 1). (**d**) Non-cathelicidin activators identified in human AMP screen activate PhoPQ when surface-displayed in KB1. Controls in which the indicated non-cathelicidin activators are surface-displayed in KB1 PhoQ H277A shown in grey. Positive (LL-37) and negative (no peptide) controls in KB1 shown in red and blue, respectively. Each plot contains data collected over *n* = 3 separate days, with results from each day shown as a separate marker. (**e**) Structures of helical non-cathelicidin human AMP activators (PDB ID – GNLY: 1L9L, NPY: 1RON). (**f**) Training and test set results for the cathelicidin sparse robust linear model. Coefficient of determination ($r^2$) and Pearson correlation coefficient ($r$) values were calculated for log10-transformed fold change and predicted fold change values.

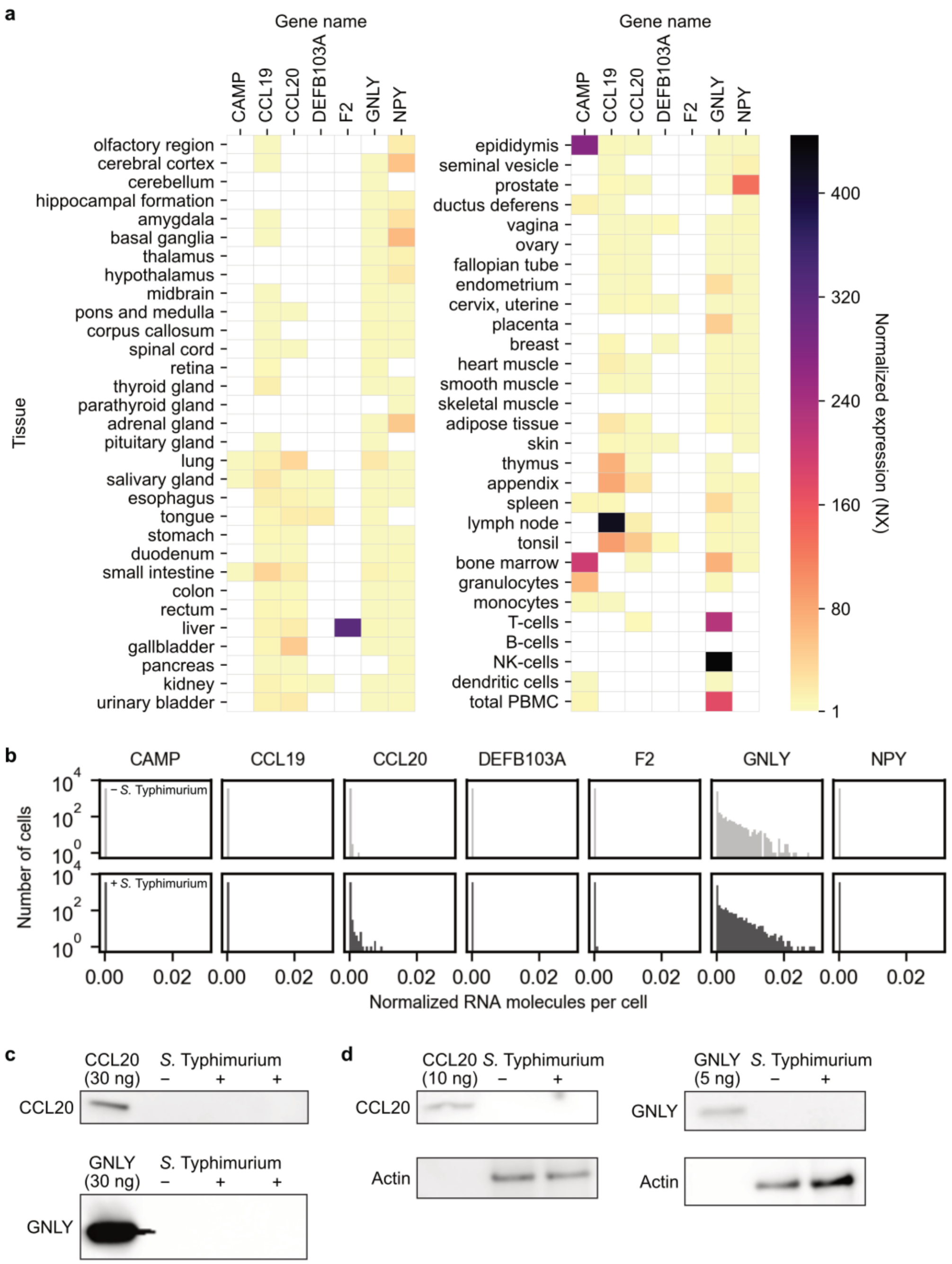


**Supplementary Figure 6.** Expression of PhoPQ-activating AMPs in human tissues.

(**a**) Tissue-specific RNA expression of the PhoPQ-activating AMPs identified in the sort-seq screen. Normalized expression (NX) values are from the Human Protein Atlas (HPA) RNA consensus tissue gene data dataset[21,22]. NX values less than 1 are indicated by white squares, consistent with the limit of detection used by HPA for classifying expression profiles. (**b**) Single-cell RNA expression in human peripheral blood mononuclear cells (PBMCs) for genes corresponding to PhoQ-activating human AMPs, as reported by Bossel Ben-Moshe *et al.* 2018[23]. RNA counts are normalized to the total number of transcripts in a given cell. Light and dark grey bars represent cells that have not and have been exposed to *S.* Typhimurium, respectively. (**c-d**) CCL20 and GNLY expression in PMA-treated THP-1 cells (**c**) and in human monocyte-derived macrophages (hMDM) (**d**) that have not (−) or have (+) been exposed to *S.* Typhimurium. The first lane is a control in which the indicated amount of purified AMP was loaded. Results in (**c**) and (**d**) are representative Western blots from $n \geq 3$ biological replicates; the two *S.* Typhimurium-exposed lanes in (**c**) are technical replicates. Bottom panels in (**d**) are actin loading controls. Where different, gene and AMP names correspond as follows: CAMP – cathelicidin, DEFB103A – hBD3, F2 – TCP.

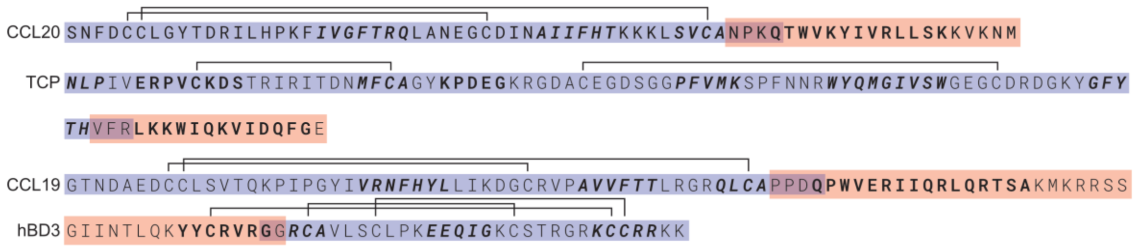


**Supplementary Figure 7.** Sequences of truncation mutants of α+β activators.

Brackets indicate disulfide bonds. α-helix and Δ α-helix fragments from **Figure 4b** are highlighted in red and blue, respectively. Bold and bold italic regions have helical and β-sheet secondary structures in the full-length peptide, respectively.

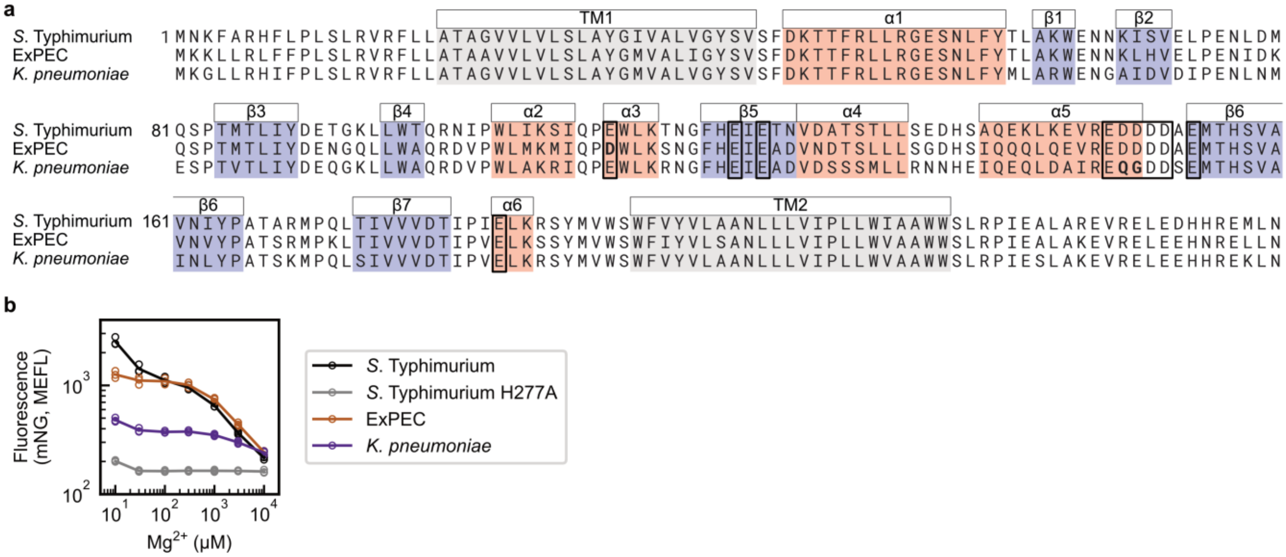


**Supplementary Figure 8.** Development of *E. coli* strains expressing ExPEC and *K. pneumoniae* PhoPQ.

(**a**) Alignment of *S.* Typhimurium, ExPEC, and *K. pneumoniae* periplasmic domains and adjacent regions. Transmembrane (TM) domains are shaded in grey and are with respect to *S.* Typhimurium. Periplasmic α-helices and β-sheets are shaded in red and blue, respectively. Boxes indicate acidic patch residues. Secondary structure and acidic patch residues are annotated according to *S.* Typhimurium as in Prost *et al.* 2008[24]. Acidic patch residues in ExPEC and *K. pneumoniae* that differ relative to *S.* Typhimurium PhoQ are bolded. (**b**) $Mg^{2+}$ dose response curve of *E. coli* strains expressing PhoP and PhoQ orthologs from *S.* Typhimurium (KB1), ExPEC (AM1), and *K. pneumoniae* (AM2), as well as a control strain expressing *S.* Typhimurium PhoP and phosphomutant PhoQ H277A (KB1 PhoQ H277A). Data was collected over $n$ = 3 separate days, with results from each day shown as a separate marker.

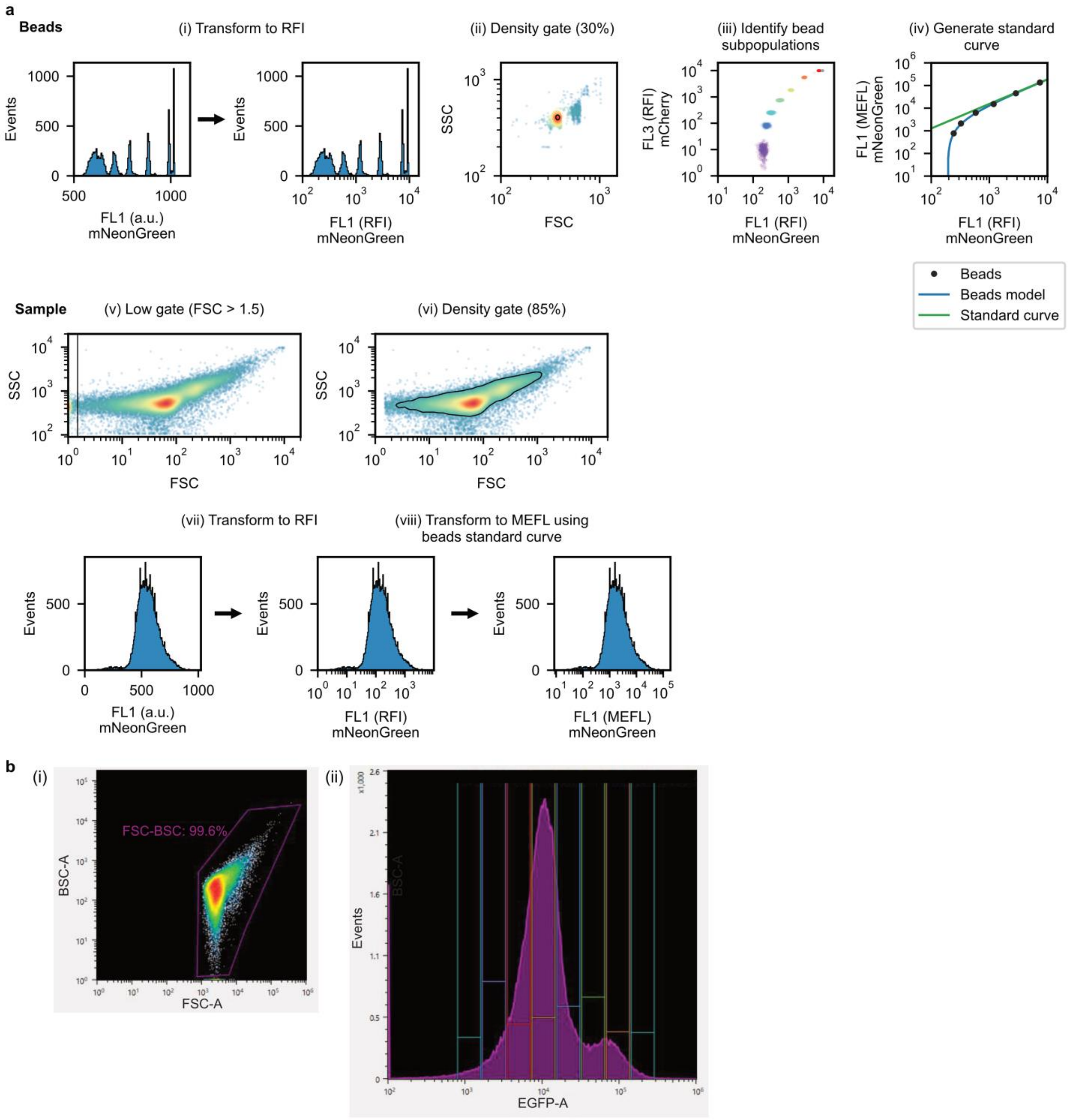


**Supplementary Figure 9.** Flow cytometry analysis and FACS gating strategies.

(**a**) For all experiments except the sort-seq experiment, all gating was performed *in silico* using FlowCal[25]. (i-iv) On each experiment day, flow cytometry data was collected for fluorescent calibration beads. (i) Beads fluorescence data was transformed from arbitrary units (a.u.) to relative fluorescence intensity (RFI). (ii) A 30% density gate was applied in forward scatter (FSC) and side scatter (SSC) to isolate beads from debris. (iii) Bead subpopulations with varying fluorescence intensities were identified using data from two fluorescence channels. (iv) RFI values were compared to manufacturer-provided molecules of equivalent fluorescein (MEFL) values for each bead subpopulation to develop a standard curve. (v-viii) Gating steps for *S.* Typhimurium and *E. coli* flow cytometry samples. (v) Samples containing *S.* Typhimurium were gated to remove events with saturating low FSC. This gating step was not applied to *E. coli* samples, where low FSC saturation was not observed. (vi) An 85% density gate was applied to remove outliers in FSC and SSC. (vii-viii) Fluorescence data was transformed from a.u. to RFI and from RFI to MEFL using the beads standard curve. (**b**) Gating strategy for cell sorting in the sort-seq experiment. (i) A gate in FSC-A (forward scatter – area) and BSC-A (back scatter – area) was hand drawn to capture a maximum number of events (approximately 99.6%) while excluding a small number of events that were likely debris. (ii) Cells were sorted into 8 hand-drawn fluorescence bins in EGFP-A (enhanced green fluorescent protein – area) that were approximately logarithmically spaced.

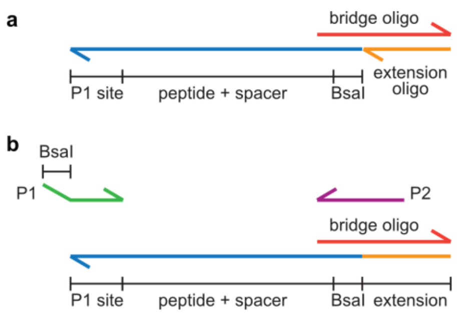


**Supplementary Figure 10.** Amplification strategy for the human AMP library oligo pool.

(**a**) Oligos in the human AMP library oligo pool (blue) were designed to contain only one primer binding site at their 3' end (P1 site, 3' end indicated by half-arrow). An additional primer binding site at the 5' end (blunt) was introduced by annealing extension (orange) and bridge (red) oligos to the oligos in the pool and performing a ligation reaction. (**b**) The resulting extended oligos were amplified via PCR. For additional details, see **Methods**.

### Supplementary Tables

**Supplementary Table 1.** Human AMP sequences, cluster assignments, basic biophysical and biochemical properties, primer sequences, and fold changes in sort-seq experiment.

**Supplementary Table 2.** Composition of sort-seq bins.

Minimum and maximum fluorescence values represent the events with the lowest and highest fluorescence values collected in each bin, as determined by indexed sort data. Percent of events is calculated based on the number of events out of 200,000 collected events with fluorescence values between the minimum and maximum values reported here.

| Bin | Percent of events (%) | Minimum fluorescence (a.u.) | Maximum fluorescence (a.u.) |
|---|---:|---:|---:|
| 1 | 1.3 | 779 | 1601 |
| 2 | 4.6 | 1659 | 3410 |
| 3 | 19.7 | 3410 | 7264 |
| 4 | 43.9 | 7264 | 14928 |
| 5 | 14.6 | 15475 | 31799 |
| 6 | 5.6 | 31801 | 65350 |
| 7 | 5.0 | 67747 | 139215 |
| 8 | 1.0 | 139220 | 296555 |

**Supplementary Table 3.** Peptide features considered for inclusion in the cathelicidin sparse robust linear model.

| Feature type | Feature | Number of descriptors | Number of features |
|---|---|---|---|
| Basic features | Charge | 1 | 1 |
| | Grand average of hydropathicity (GRAVY) | 1 | 1 |
| | Isoelectric point | 1 | 1 |
| | Length | 1 | 1 |
| | Molecular weight | 1 | 1 |
| | Aromaticity | 1 | 1 |
| | Instability | 1 | 1 |
| Residue composition | Amino acid composition | 1 | 20 |
| | Dipeptide composition | 1 | 400 |
| | Pairwise relative amino acid composition | 1 | 380 |
| Autocorrelation | Normalized Moreau-Broto autocorrelation | 8 | $8 \times \delta_{max}$ |
| | Moran autocorrelation | 8 | $8 \times \delta_{max}$ |
| | Geary autocorrelation | 8 | $8 \times \delta_{max}$ |
| Physicochemical composition | Composition | 7 | 21 |
| | Transition | 7 | 21 |
| | Distribution | 7 | 105 |
| Sequence order features | Sequence-order-coupling number | 2 | $2 \times \delta_{max}$ |
| | Pseudo-amino acid composition (PAAC) | 1 | $20 + \lambda_{max}$ |
| | Amphiphilic PAAC (APAAC) | 2 | $20 + 2 \times \lambda_{max}$ |

**Supplementary Table 4.** Features used in final sparse robust linear model for cathelicidin-like peptides.

| Feature Name | Interpretation | Coefficient in Model | Feature Importance |
|---|---|---|---|
| MoranAuto_Hydrophobicity7 | Moran autocorrelation measuring pairwise correlation in hydrophobicity for residues separated by δ = 7 positions | 0.212 | 0.094 |
| PAAC29 | Pseudo-amino acid composition measuring pairwise correlations in hydrophobicity, hydrophilicity, and side chain mass for residues separated by δ = 9 positions | 0.047 | 0.093 |
| pc(G,R) | Relative fraction of G residues to R residues | -0.309 | 0.089 |
| Charge | Net charge of the peptide primary amino acid sequence | 0.029 | 0.087 |
| pc(T,K) | Relative fraction of T residues to K residues | -0.298 | 0.087 |
| _HydrophobicityD2001 | Fraction of peptide length containing the first residue with neutral hydrophobicity (G, A, S, T, P, H, or Y) | -0.003 | 0.082 |
| MoreauBrotoAuto_Hydrophobicity10 | Normalized Moreau-Broto autocorrelation measuring pairwise correlation in hydrophobicity for residues separated by δ = 10 positions | -0.230 | 0.074 |
| KD | Fraction of KD dipeptides | 0.026 | 0.072 |
| GearyAuto_Hydrophobicity7 | Geary autocorrelation measuring pairwise correlation in hydrophobicity for residues separated by δ = 7 positions | -0.169 | 0.069 |
| PAAC28 | Pseudo-amino acid composition measuring pairwise correlations in hydrophobicity, hydrophilicity, and side chain mass for residues separated by δ = 8 positions | 0.053 | 0.061 |
| GearyAuto_ResidueASA8 | Geary autocorrelation measuring pairwise correlation in solvent-accessible surface area for residues separated by δ = 8 positions | -0.215 | 0.052 |
| TS | Fraction of TS dipeptides | -0.003 | 0.050 |
| GearyAuto_Mutability3 | Geary autocorrelation measuring pairwise correlation in mutability for residues separated by δ = 3 positions | 0.258 | 0.048 |
| RC | Fraction of RC dipeptides | 0.229 | 0.042 |
| Constant | n/a | -0.091 | n/a |

**Supplementary Table 5.** Bacterial strains used in this study.

Only base strains are listed; derivatives of these strains were also generated by transforming these strains with plasmids listed in **Supplementary Table 6**.

| **Strain description** | **Source** | **Identifier** |
|---|---|---|
| *Salmonella enterica* subsp. *enterica* serovar Typhimurium strain 14028 | ATCC | ATCC 14028 |
| *Salmonella enterica* subsp. *enterica* serovar Typhimurium 14028s Δ*phoQ* | BEI Resources | NR-40554 |
| *E. coli* BW30007 | Coli Genetic Stock Center (from Zhou et al. 2003[26]) | CGSC# 7942 |
| *E. coli* BW30007 carrying *S.* Typhimurium *phoP* and *phoQ* | This paper | KB1 |
| *E. coli* BW30007 carrying *S.* Typhimurium *phoP* and *phoQ* H277A | This paper | KB1 PhoQ H277A |
| *E. coli* BW30007 carrying ExPEC (EC958 (ST131)) *phoP* and *phoQ* | This paper | AM1 |
| *E. coli* BW30007 carrying *K. pneumoniae* (Kp52145) *phoP* and *phoQ* | This paper | AM2 |

**Supplementary Table 6.** Plasmids used in this study.

| Name | Contents | Source | Addgene ID |
|---|---|---|---|
| pAM002 | $P_{pmrD}$-sfGFP reporter, PhoP overexpression | This paper | 170006 |
| pAM003 | $P_{mgtC}$-sfGFP reporter, PhoP overexpression | This paper | 170007 |
| pAM004 | $P_{ugtL}$-sfGFP reporter, PhoP overexpression | This paper | 170008 |
| pAM005 | $P_{virK}$-sfGFP reporter, PhoP overexpression | This paper | 170009 |
| pAM006 | $P_{pagC}$-sfGFP reporter, PhoP overexpression | This paper | 170010 |
| pAM007 | $P_{slyB}$-sfGFP reporter, PhoP overexpression | This paper | 170011 |
| pKB222 | $P_{virK}$-mNG reporter, LL-37 peptide display | This paper | 170012 |
| pKB223 | $P_{virK}$-mNG reporter, C18G peptide display | This paper | 170013 |
| pKB224 | $P_{virK}$-mNG reporter, NC peptide display | This paper | 170014 |
| pKB225 | $P_{virK}$-mNG reporter, no displayed peptide | This paper | 170015 |
| pKB226 | $P_{virK}$-mNG reporter, defensin HNP-1 peptide display | This paper | 170016 |
| pKB230 | $P_{virK}$-mNG reporter, protegrin-1 peptide display | This paper | 170017 |
| pKB231 | $P_{virK}$-mNG reporter, cecropin P1 peptide display | This paper | 170018 |
| pKB232 | $P_{virK}$-mNG reporter, murine CRAMP peptide display | This paper | 170019 |
| pKB233 | $P_{virK}$-mNG reporter | This paper | 170040 |
| pKB234 | $P_{virK}$-mNG reporter, hBD3 peptide display | This paper | 170021 |
| pKB234.1 | $P_{virK}$-mNG reporter, hBD3 α-helix peptide display | This paper | 170022 |
| pKB234.2 | $P_{virK}$-mNG reporter, hBD3 Δ α-helix peptide display | This paper | 170023 |
| pKB235 | $P_{virK}$-mNG reporter, CCL20 peptide display | This paper | 170024 |
| pKB235.2 | $P_{virK}$-mNG reporter, CCL20 α-helix peptide display | This paper | 170025 |
| pKB235.4 | $P_{virK}$-mNG reporter, CCL20 Δ α-helix peptide display | This paper | 170026 |
| pKB236 | $P_{virK}$-mNG reporter, TCP peptide display | This paper | 170027 |
| pKB236.1 | $P_{virK}$-mNG reporter, TCP Δ α-helix peptide display | This paper | 170028 |
| pKB236.2 | $P_{virK}$-mNG reporter, TCP α-helix peptide display | This paper | 170029 |
| pKB237 | $P_{virK}$-mNG reporter, CCL19 peptide display | This paper | 170030 |
| pKB237.1 | $P_{virK}$-mNG reporter, CCL19 Δ α-helix peptide display | This paper | 170031 |
| pKB237.2 | $P_{virK}$-mNG reporter, CCL19 α-helix peptide display | This paper | 170032 |
| pKB238 | $P_{virK}$-mNG reporter, NPY peptide display | This paper | 170033 |
| pKB239 | $P_{virK}$-mNG reporter, GNLY peptide display | This paper | 170034 |
| pKB252 | $P_{virK}$-mNG reporter, K6L9 peptide display | This paper | 170035 |
| pKB253 | $P_{virK}$-mNG reporter, K5L7 peptide display | This paper | 170036 |
| pKB254 | $P_{virK}$-mNG reporter, H6L9 peptide display | This paper | 170037 |
| pKB255 | $P_{virK}$-mNG reporter, Melittin peptide display | This paper | 170038 |
| pKB256 | $P_{virK}$-mNG reporter, Bac2a peptide display | This paper | 170039 |

**Supplementary Table 7.** Commercial sources of peptides used in this study.

| Peptide | Product name | Manufacturer | Catalog number |
|---|---|---|---|
| CCL19 | MIP-3β Human E. coli | BioVendor | Cat# RBG10228100 |
| CCL20 | Recombinant Human MIP-3α / CCL20 | Cell Sciences | Cat# CRH317B |
| GNLY | Granulysin Human E. coli | BioVendor | Cat# RD172327100 |
| hBD3 | hBD-3, β-Defensin-3, human | Anaspec | Cat# AS-60741 |
| LL-37 | LL-37, Antimicrobial Peptide, human | Anaspec | Cat# AS-61302 |
| NPY | Neuropeptide Y (human, rat) | Tocris | Cat# 1153 |
| TCP C-terminal α-helix | N/A (custom synthesis) | Genscript | N/A |

**Supplementary Table 8.** Primers used to generate next-generation sequencing amplicons.

Partial adapter sequences required for Genewiz Amplicon-EZ are capitalized and underlined. Sample barcodes are capitalized with no underline and primer binding sites are in lower case.

| Sample | Primer 1 | Primer 1 sequence | Primer 2 | Primer 2 sequence |
|---|---|---|---|---|
| Pre-sort | KB640 | GACTGGAGTTCAGACGTGTGCTCTTCCGATCTttaggcttcggcatggggtcaggtg | KB585 | ACACTCTTTCCCTACACGACGCTCTTCCGATCTagctgcgggtatcggagga |
| Bin 1 | KB649 | TGCCTTGATCttcggcatggggtcaggtg | KB659 | TGCCTTGATCagctgcgggtatcggagga |
| Bin 2 | KB642 | CGCTCAGTTCttcggcatggggtcaggtg | KB652 | CGCTCAGTTCagctgcgggtatcggagga |
| Bin 3 | KB650 | CTACTCAGTCttcggcatggggtcaggtg | KB660 | CTACTCAGTCagctgcgggtatcggagga |
| Bin 4 | KB644 | ATATGAGACGttcggcatggggtcaggtg | KB654 | ATATGAGACGagctgcgggtatcggagga |
| Bin 5 | KB645 | CTTATGGAATttcggcatggggtcaggtg | KB655 | CTTATGGAATagctgcgggtatcggagga |
| Bin 6 | KB646 | TAATCTCGTCttcggcatggggtcaggtg | KB656 | TAATCTCGTCagctgcgggtatcggagga |
| Bin 7 | KB647 | GCGCGATGTTttcggcatggggtcaggtg | KB657 | GCGCGATGTTagctgcgggtatcggagga |
| Bin 8 | KB648 | AGAGCACTAGttcggcatggggtcaggtg | KB658 | AGAGCACTAGagctgcgggtatcggagga |
| No gate | KB651 | TCGTCTGACTttcggcatggggtcaggtg | KB661 | TCGTCTGACTagctgcgggtatcggagga |
